# Extracellular vesicles produced by the human commensal gut bacterium *Bacteroides thetaiotaomicron* affect host immune pathways in a cell-type specific manner that are altered in inflammatory bowel disease

**DOI:** 10.1101/2021.03.20.436262

**Authors:** Lejla Gul, Dezso Modos, Sonia Fonseca, Matthew Madgwick, John P. Thomas, Padhmanand Sudhakar, Régis Stentz, Simon R. Carding, Tamas Korcsmaros

**Affiliations:** Earlham Institute, Norwich, UK; Gut Microbes and Health Research Programme, Quadram Institute Bioscience Norwich, UK; Department of Gastroenterology, Norfolk and Norwich University Hospital, Norwich, UK; KU Leuven Department of Chronic Diseases, Metabolism and Ageing, Translational Research Center for Gastrointestinal Disorders (TARGID), Leuven, Belgium; Norwich Medical School, University of East Anglia, Norwich, UK

**Keywords:** extracellular vesicles, host-microbe interactions, single-cell data analysis, ulcerative colitis, Toll-like receptor pathway

## Abstract

The gastrointestinal (GI) tract is inhabited by a complex microbial community, which contributes to its homeostasis. Disrupted microbiome can cause GI-related diseases, including inflammatory bowel disease, therefore identifying host-microbe interactions is crucial for better understanding gut health. Bacterial extracellular vesicles (BEVs), released into the gut lumen, can cross the mucus layer and access underlying immune cells. To study cross-kingdom communication between BEVs and host, we focused on the influence of BEVs, generated by *Bacteroides thetaiotaomicron* (VPI-5482), on host immune cells. Using single-cell RNA sequencing data and host-microbe protein-protein interaction networks, we examined the potential effect of BEVs on dendritic cells, macrophages and monocytes with particular focus on the Toll-like receptor (TLR) pathway. We identified biological processes affected in each immune cell type, and also cell-type specific processes (e.g myeloid cell differentiation). The TLR pathway analysis highlighted that BEV targets differ among cells and even between the same cells in healthy versus disease (ulcerative colitis) conditions. Our *in silico* findings were validated in BEV-monocyte co-cultures demonstrating the requirement for TLR4 in BEV-elicited NF-κ B activation. This study demonstrates that both cell-type and health condition influence BEV-host communication. The results and the pipeline can facilitate BEV-based therapy development for the treatment of IBD.

## Introduction

The human gastrointestinal (GI) tract microbiota consisting of bacteria, viruses, archaea, and eukaryotic microbes, contributes to intestinal homeostasis by communicating with various host cells in the intestinal mucosa. Structural, compositional and functional alterations of the microbiota (“dysbiosis”) are associated with various GI-related diseases, including Crohn’s disease (CD) and ulcerative colitis (UC), two major forms of inflammatory bowel disease (IBD) [1]. Dysbiosis in IBD is characterised by a reduction in bacterial diversity (UC) or altered composition (CD) that involves *Bacteroides* and *Firmicutes* species [2]. Despite recent advances in our understanding of IBD pathogenesis, the complex interactions between the dysbiotic gut microbiota and the host mucosa that result in aberrant immune activation and inflammation in the gut, are yet to be defined in detail.

*Bacteroides thetaiotaomicron* (Bt) is a Gram-negative anaerobe that is a major constituent of the human caecal and colonic microbiota [3]. The administration of Bt in murine models of IBD ameliorates inflammation [4, 5] with the anti-inflammatory effects being at least in part mediated by its production of bacterial extracellular vesicles (BEVs). BEVs are released by both commensal Gram-negative and Gram-positive bacteria and have the potential to mediate cross-kingdom interactions with host cells via the delivery of their contents and cargo to affect host cell physiology and function [4]. BEVs produced by Gram-negative bacteria such as Bt (outer membrane vesicles; OMVs) are small, spherical bilayered structures (20-400 nm) composed of phospholipids, lipopolysaccharides, peptidoglycan, outer membrane proteins, periplasmic contents including proteins, and some inner membrane and cytoplasmic fractions [6]. BEVs can permeate through the sterile mucus layer of the colon to access and transmigrate boundary intestinal epithelial cells through different routes [7] enabling them to interact with underlying mucosal immune cells [8–12] and the intestinal vasculature which facilitates their wider, systemic dissemination [7, 12, 13].

BEVs contain various microbe associated molecular patterns (MAMPs) enabling them to interact with immune cells expressing appropriate pattern recognition molecules (PRRs) such as Toll-like receptors (TLR). BEVs can trigger TLR signalling in various immune cells including dendritic cells (DC) [12] resulting in the activation of different transcription factors, and depending on the cell-type and TLRs involved, generate diverse immune effector cell outcomes [14]. These effects may potentially be anti-inflammatory and BEVs have been incorporated into probiotic-based therapeutics in murine models of IBD [4, 5]. Despite the advent of biologic therapies in IBD, ∼25% of patients with UC and up to 75% with CD eventually require surgical intervention, highlighting the need for novel therapeutic strategies. One such strategy that is being actively explored is the ability to modulate the host immune system through microbiota-based therapies [15]. Given the ability of Bt BEVs to influence host immune cell signalling they may have untapped therapeutic potential.

However, the effects of Bt BEVs on different host immune cells are poorly understood. Single-cell transcriptomics (scRNAseq) provides an opportunity to understand how Bt BEVs can influence gut mucosal immune cell populations with cell-type specific resolution. Of particular interest are monocytes, macrophages and DCs, which play key roles in initiating and determining the outcome of local and systemic immune responses to non-harmful and harmful stimuli [16], and shaping the immune response in IBD [17].

To better define the molecular basis of the interactions between Bt BEVs and host immune cells we have utilised RNAseq datasets to study the effect of Bt BEV proteins on host signalling pathways in monocytes, macrophages, and DCs. This workflow enables comparisons of the effect of BEVs on immune cells at different stages of their development, under healthy versus disease (in this case UC) conditions. In a proof-of-concept example we experimentally demonstrate the potential impact of Bt BEVs on the TLR4 pathway in the healthy GI-tract and in UC patients.

## Material & Methods

### Characterisation of Bt BEV proteins

Identification of BEV proteins derived from Bt is described in detail in [18]. Briefly, a comparative proteomic analysis of BEVs generated either in vitro by Bt cultured in a complex medium or in vivo by isolating BEVs from the caecal content of Bt mono-colonized germ-free mice identified a total of 2068 proteins. BEVs were concentrated by crossflow filtration, separated by size exclusion chromatography and recovered by ultracentrifugation followed by protein extraction. Differential expressions of BEV proteins obtained under the two conditions were explored using tandem mass tagging (TMT) combined with liquid chromatography mass spectrometry (LC-MS/MS). Data sets were analysed using the Proteome Discoverer v2.1 software.

### Single-cell transcriptomic datasets analysis

A publicly available scRNAseq dataset describing gene expressions in 51 cell-types from the colon in three conditions (healthy, non-inflamed UC, and inflamed UC) was analysed by using the average expression of genes [19]. To discard genes expressed at extremely low levels, we applied a z-score test based on the method of Hart *et al* [20]. A gene was considered not to be expressed if its log2 expression value was less than 2 standard deviations of the mean expressed genes in that cell. From the 51 cell-type datasets, cycling monocytes, inflammatory monocytes, macrophages, DC1 (healthy mucosa-related subset) and DC2 (inflammation-related subset) populations appearing in healthy and non-inflamed UC conditions were selected for further analysis. While the original dataset contains inflamed samples as well, to avoid inflammation-related bias in cell communication, we focused our analysis on non-inflamed samples from the same UC patients.

### THP-1 monocyte transcriptomic analysis

Two publicly available bulk RNAseq datasets of the human monocytic cell line THP-1 were used for experimental validation. Raw count from GSE132408 and GSE157052 datasets were normalized using the DESeq2 package in R. Due to the different gene symbols and gene IDs in the datasets, we unified them to gene symbols using Uniprot. Applying the same protocol as for the single-cell data, selection of expressed genes by z-score test and log2 transformation were used.

### Constructing a host cell-BEV interactome

We predicted the effect of BEV proteins on different cell-types based on host-microbe protein-protein interaction (PPI) networks using our MicrobioLink pipeline [21]. The connections were based on experimentally verified domain-motif interactions from the Eukaryotic Linear Motif (ELM) database [22]. It was assumed that a BEV protein containing a domain can bind to a human protein having the corresponding interacting motif within its sequence. First, we downloaded the sequence of BEV and human proteins from the Uniprot database [23]. Then the domains of BEV proteins were predicted by InterProScan and human motifs identified by the ELM database. To avoid large numbers of false-positive PPIs, a quality filter was applied using IUPred tool [24] which uses scores based on two methods (IUPred and ANCHOR2) to measure residue-level energy terms. The energy terms correlate how intrinsically disordered the protein region is. Higher disordered regions are more accessible for the bacterial domain. Two cut-off values (IUPred > 0.5 and ANCHOR2 > 0.4) were set up to select human motifs which are presented out of globular domains and at an intrinsic disordered protein region.

### Functional analysis of BEV target proteins

Functional analysis was performed using the Gene Ontology (GO) database. GO database orders the annotations in a tree-like structure where parent and child categories are represented in a hierarchical way. GOrilla was used to highlight the enriched biological processes of the BEV targets in different cell-types [25]. As a background dataset, all expressed genes were examined in cells facilitating the identification of cell-type specific functions. We used REVIGO to reduce the dimensionality of the annotations, thereby avoiding the overlapping processes that belong to the same function and identify significant differences among functions [26]. simRel scores were applied to measure the GO semantic similarity. To visualise the functional overlap among cell-types, InteractiVenn was used [27]. Although this analysis is suitable for depicting processes that are specific to a cell-type or condition due to the large number of BEV interacting proteins in each cell-type, the output of this analysis focuses mainly on common processes. A more fine-grained analysis can be achieved by involving gene expression values, and not only the presence or absence of a gene’s expression when establishing condition specific differences.

### Cell-type and condition specific TLR pathway modelling

To perform cell-type and condition specific analyses we utilised a different approach. Members of the TLR pathway were derived from the Reactome database [28] and a list of proteins involved in the signalling and interactions between them were obtained from OmniPath database [29]. Signalling in different cell-types was interpreted by adding the expression values from scRNAseq datasets (monocytes, dendritic cells, macrophage) and bulk RNAseq (THP-1 cells). To compare the signal flow under different conditions (healthy and non-inflamed UC), expression values were added to the genes/proteins.

### Isolation and characterisation of Bt BEVs for the experimental validation

Bt (strain VPI-5482) was cultured in 500 mL of Bacteroides Defined Media plus (BDM+) (Supplementary Table 1) at 37°C in an anaerobic cabinet. Cells were harvested after 17.5 h at an OD_600nm_ of 1.98. BEVs were isolated following a method adapted from Stentz et al [30]. Briefly, cultures (500 mL) were centrifuged at 6000 g for 1h at 4°C and the supernatants vacuum-filtered through polyethersulfone (PES) membranes (0.22 μ pore-size) (Sartorius) to remove debris and cells. Supernatants were concentrated by ultrafiltration at 100 kDa molecular weight cut-off, (Vivaspin 50R, Sartorius), the retentate was rinsed with 500 mL of PBS (pH 7.4) and concentrated to 5 mL. Any precipitate was spun down at 15000 g for 20min and the supernatant filter sterilised (0.22 μm pore size). The final BEVs suspensions were stored at 4°C.

The size and concentration of the isolated BEVs was determined using a ZetaView PMX-220 TWIN instrument according to manufacturer’s instructions (Particle Metrix GmbH). Aliquots of BEVs suspension were diluted 1000- to 20,000-fold in particle-free PBS for analysis. Size distribution video data was acquired using the following settings: temperature: 25°C; frames: 60; duration: 2 seconds; cycles: 2; positions: 11; camera sensitivity: 80 and shutter value: 100. The ZetaView NTA software (version 8.05.12) was used with the following post acquisition settings: minimum brightness: 20; max area: 2000; min area: 5 and trace length: 30.

### TLR-signalling in THP1-Blue cells

THP1-Blue NF-κ B reporter cell line (Invivogen) was derived from the human THP-1 monocytic cell line by stable integration of an NF-B-inducible secreted alkaline phosphatase κ (SEAP) reporter construct. THP1-Blue cells were cultivated in RPMI-1640 (Sigma-Aldrich) supplemented with 10% heat-inactivated FBS (Biosera), 1% Pen/Strep (Sigma-Aldrich) and 100 µg/mL Normocin (Invivogen) at 37°C and 5% CO_2_ in a humidified incubator. To maintain selection pressure during cell subculturing, 10 µg/mL blasticidin (Invivogen) was added to the growth medium at every other passage. To confirm TLR4 mediated activation THP-1 cells were seeded in flat-bottomed 96-well plates at a density of 5×10^5^ cells/mL and incubated with *E. coli* derived LPS (10 ng/mL, Sigma-Aldrich) 1 h at 37°C. Control cultures contained complete media only. In some cases, cells were pre-treated with the TLR4 inhibitor CLI-095 (2 µg/mL) (Invivogen) and incubated for 1.5 h at 37°C and 5% CO_2_ in a humidified incubator. For BEV-THP-1 co-culture cells were incubated for 24 h with different concentrations of BEVs (3×10^9^, 3×10^8^, 3×10^7^, and 3×10^6^/mL) after which 20 µL of the cell suspension was added to flat-bottomed 96-well plates, mixed with 180 µL of Quanti-Blue (Invivogen) colorimetric assay reagent and incubated for 1 h at 37°C. Secreted alkaline phosphatase (SEAP) levels were quantified by absorbance reading at 620 nm. All incubations were performed in triplicate.

### Statistical analysis

Data were subjected to one-way ANOVA followed by Dunnett’s multiple comparison post hoc test using GraphPad Prism 5 software. Statistically significant differences between two mean values were established by adjusted p-value < 0.05. Data are presented as the mean ± standard deviation.

### Data availability

Raw scRNAseq data was extracted from Smillie et al [19]. Bulk transcriptomics for THP-1 cell line analysis can be found in GEO [GSE132408, GSE157052]. The workflow containing Python and R scripts, input files and results is accessible on GitHub (https://github.com/korcsmarosgroup/BT_BEV_project/).

## Results

### The BEV - immune cell protein interactome

To analyse the effect of BEV proteins on human cell-type specific signalling pathways we developed a computational workflow to process single-cell data, combine information from network resources, and incorporate bioinformatics prediction tools [Figure 1].

**Figure 1:**
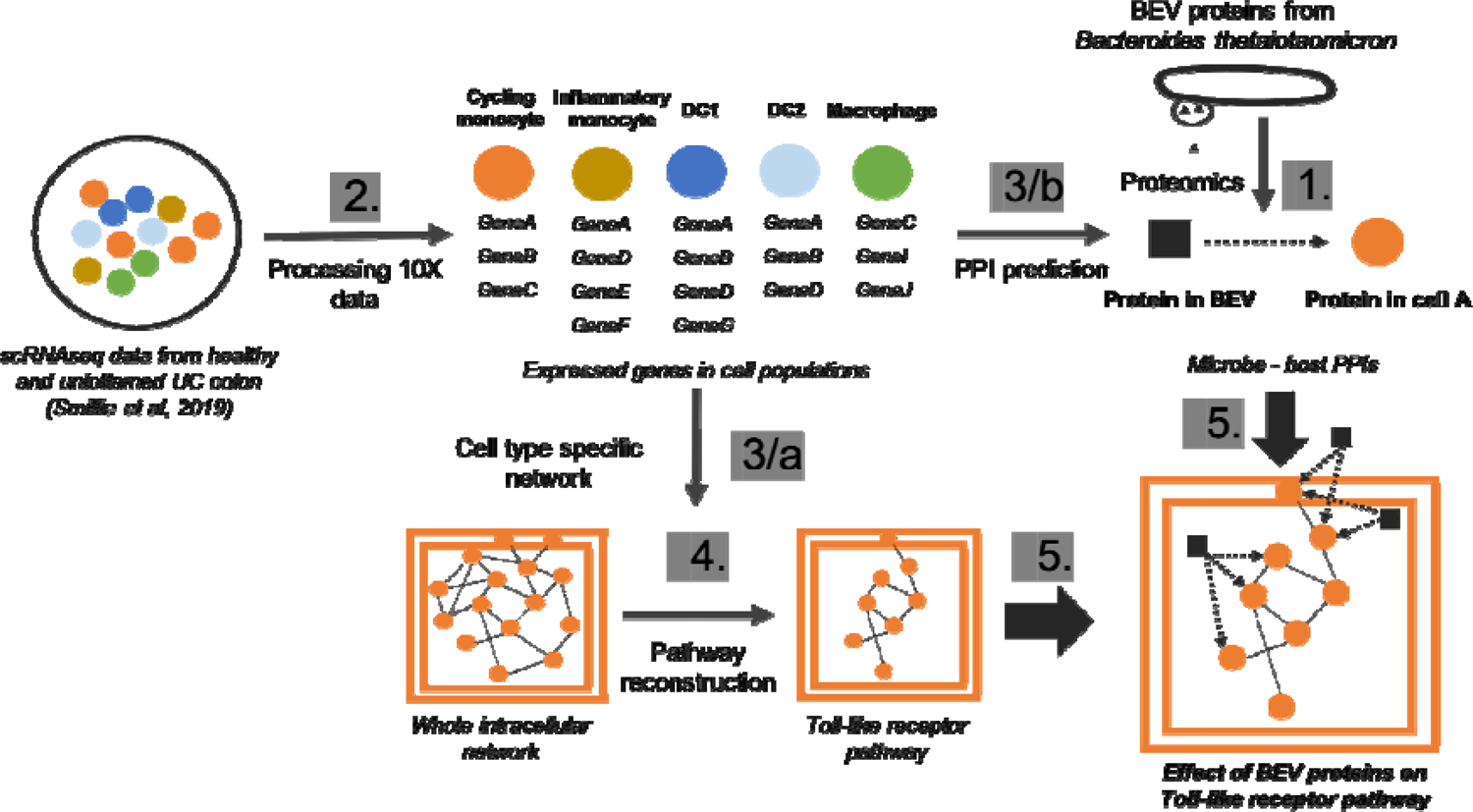
Computational workflow to analyse cell-type specific effects of BEVs. Numbers indicate the sequence of the main steps: 1, Extraction of BEV proteins from the proteomic dataset 2, Processing the raw single-cell transcriptomics from human colon 3/a, Creating cell-type specific network using protein-protein interactions from OmniPath [29] 3/b, Predicting protein-protein interactions (PPIs) between BEV and host proteins in each cell-type separately 4, Reconstruction of Toll-like receptor pathway using Reactome database [28] 5, Combining cell-specific signalling with BEV targeted human proteins

[FIG]

Using this workflow, we identified potential candidates from the proteome of BEVs obtained from the caecum of germ-free mice monocolonized with Bt, which totalled 2068 proteins [18]. For host cells scRNAseq data identified genes expressed in each of five immune cell-types [Figure 2]. For the purpose of developing the protein-protein interaction (PPI) network, we assumed that all of the expressed genes were translated into functional proteins.

**Figure 2:**
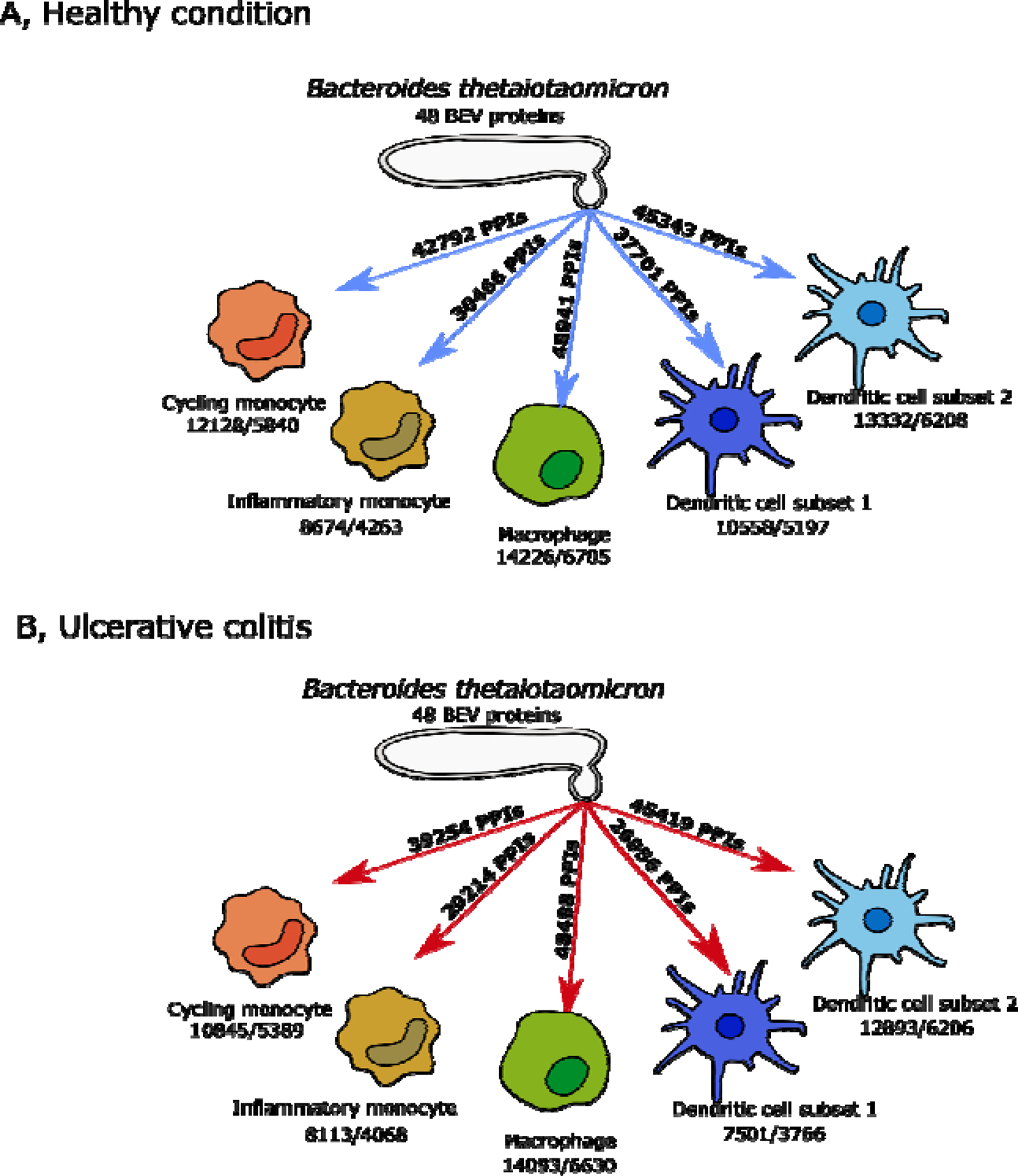
Interactions of 48 BEV proteins with monocytes, macrophages and dendritic cells in healthy (A) and UC (B) conditions. Number of expressed genes/number of interacting proteins are highlighted for each cell-type.<colcnt=1>

BEVs can interact with the host *via* cell surface receptors and after internalisation, with cytoplasmic receptors. We did not therefore filter host proteins based on their cellular location. Despite the large number of BEV-human PPIs [Figure 2] the majority of bacterial proteins were hubs indicating they can potentially interact with thousands of host proteins. The majority of these bacterial proteins are hydrolases, proteases, and other catabolic enzymes without a specific cleavage site. In terms of individual interactions, five BEV proteins [Table 1] target the same host protein PAPD5, a non-canonical poly(A) polymerase [31].

**Table 1:**
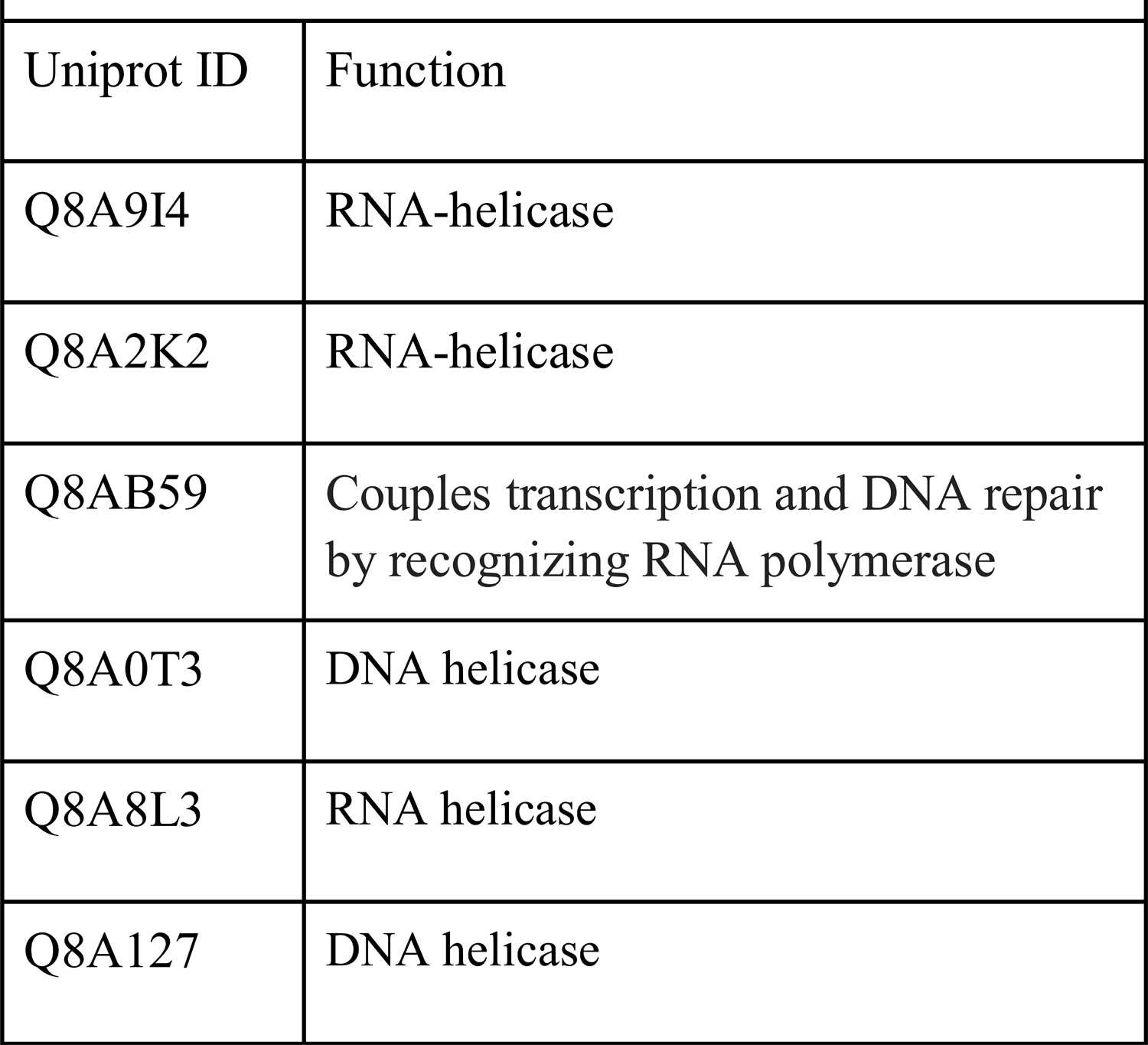
Function of BEV proteins targeting PAPD5

### Functional analysis

Cell-type specific BEV-host interactomes are complex due to the large number of proteins and interactions involved. Therefore, a functional analysis based on the GO database was initially carried out to identify the biological processes affected by microbial proteins in healthy (non-inflamed) and inflamed UC conditions. Most of the over-represented functions were overlapping among the different cell-types. However, comparing the cells under different conditions enabled us to identify specific effects of BEV proteins with the unique functions [Supplementary Table 2-3, Supplementary Figure 1-3].

**Figure 3:**
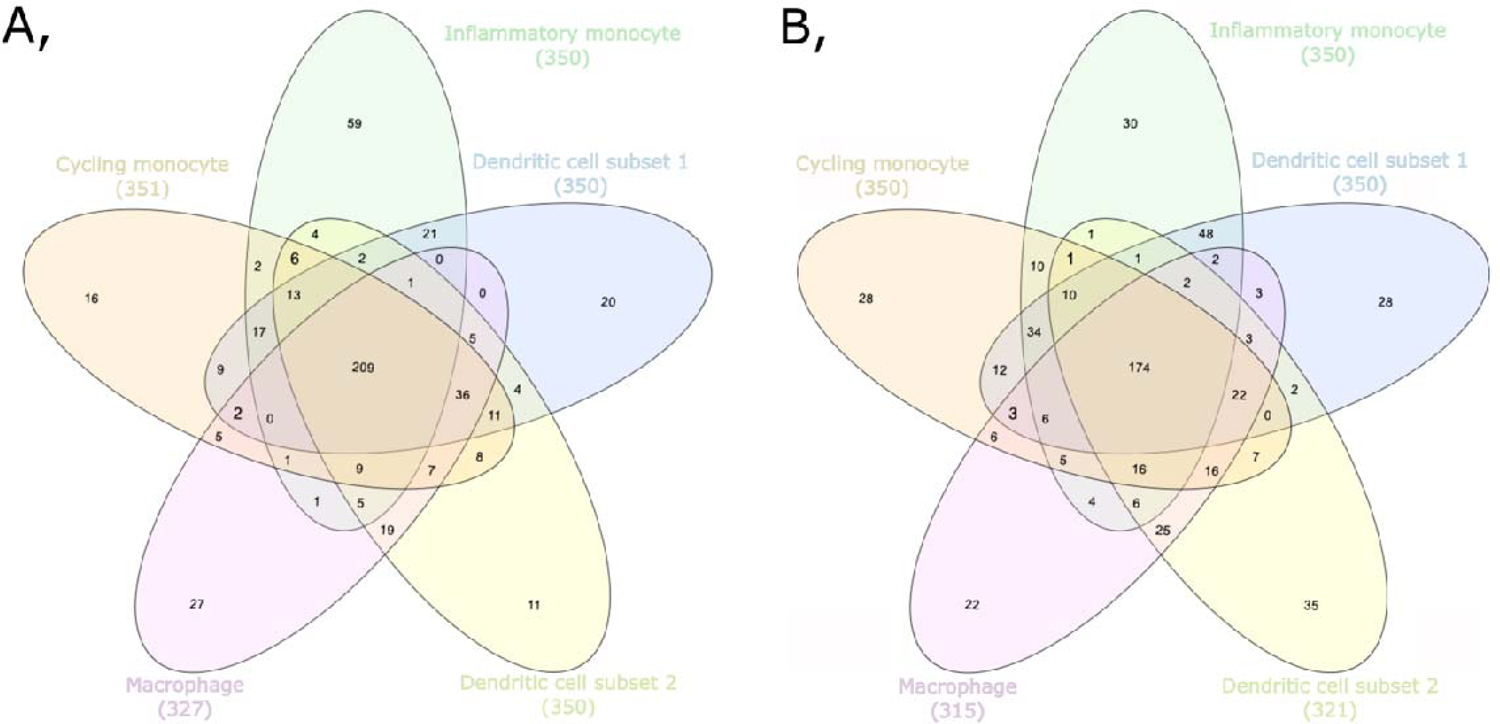
Overlap of biological processes over-represented in the BEV-host interactomes corresponding to cell-types in healthy (A) and uninflamed UC (B) conditions

In the healthy state 209 functions were shared among the five cell-types containing basic cellular functions, such as chromatin organisation and macromolecule synthesis. Most of the unique processes (59) were found in inflammatory monocytes and were related to the endoplasmic reticulum (ER), apoptosis and myeloid cell differentiation. Counter to these results, in cycling monocytes - in terms of unique functions (16) - cell cycle-related processes were uncovered. Interestingly, among BEV targets in DC1 cells (20) somatic diversification of immune receptors and B cell apoptosis were uniquely over-represented. In contrast, negative regulation of myeloid leukocyte mediated immunity and cell differentiation were prominent in DC2 cells (11). Among BEV-targeted human proteins, the signalling pathways of both the epidermal growth factor (EGF) receptor and the regulation of transforming growth factor beta (TGF-beta) receptor were affected specifically in macrophages, based on 27 individual processes [Figure 3/A].

BEV targets in the non-inflamed UC state included 174 overlapping processes that play vital roles in cell function. Uniquely over-represented functions were observed in inflammatory monocytes (30) that were similar in non-inflamed UC and healthy conditions and included positive regulation of the endoplasmic-reticulum-associated protein degradation (ERAD) pathway and intrinsic apoptotic signalling pathways. Among the 28 cycling monocytes-related annotations, similarly to the healthy condition, the cell cycle associated proteins were overrepresented. Here, we also found the negative regulation of G1/S phase transition overrepresented. Other targeted human proteins identified in this study are involved in the regulation of DNA repair and cyclin-dependent protein kinase activity, positive regulation of protein ubiquitination, and signal transduction by p53 class mediator. Whereas BEV proteins affected cell-cycle processes in DC2 (35), target proteins in DC1 (28) related to vesicle fusion, negative regulation of apoptotic signalling pathways, and the intracellular steroid hormone receptor signalling pathway. Among the 22 unique processes in macrophages, regulation of RAS protein signal transduction, base-excision repair, and diverse histone modification steps were identified [Figure 3/B].

In both conditions, macromolecule metabolism, DNA-related processes, and RNA-related processes were affected in all five cell-types by BEVs. Additionally, endoplasmic reticulum (ER)-stress response related processes and vesicle organisation and transport were influenced by BEVs in most cell-types.

### Effect of BEV proteins on TLR pathway in dendritic cells, monocytes and macrophages under healthy and UC conditions

As previous results established that Bt may alter immune pathways, we focused on the potential interactions between BEVs and TLR pathways. To do this, we created cell-type and condition specific signalling networks for BEVs and TLR pathways based on the scRNAseq data. These networks revealed that whilst the expression of TLR pathway-related transcription factors remained the same in both healthy and non-inflamed UC conditions in all examined cell-types, the level of TLR receptor expression was different amongst different immune cell-types. Due to the cell-type specific expression of different pathway members, BEV proteins established diverse interactions with immune cells [Figure 4-7].

**Figure 4:**
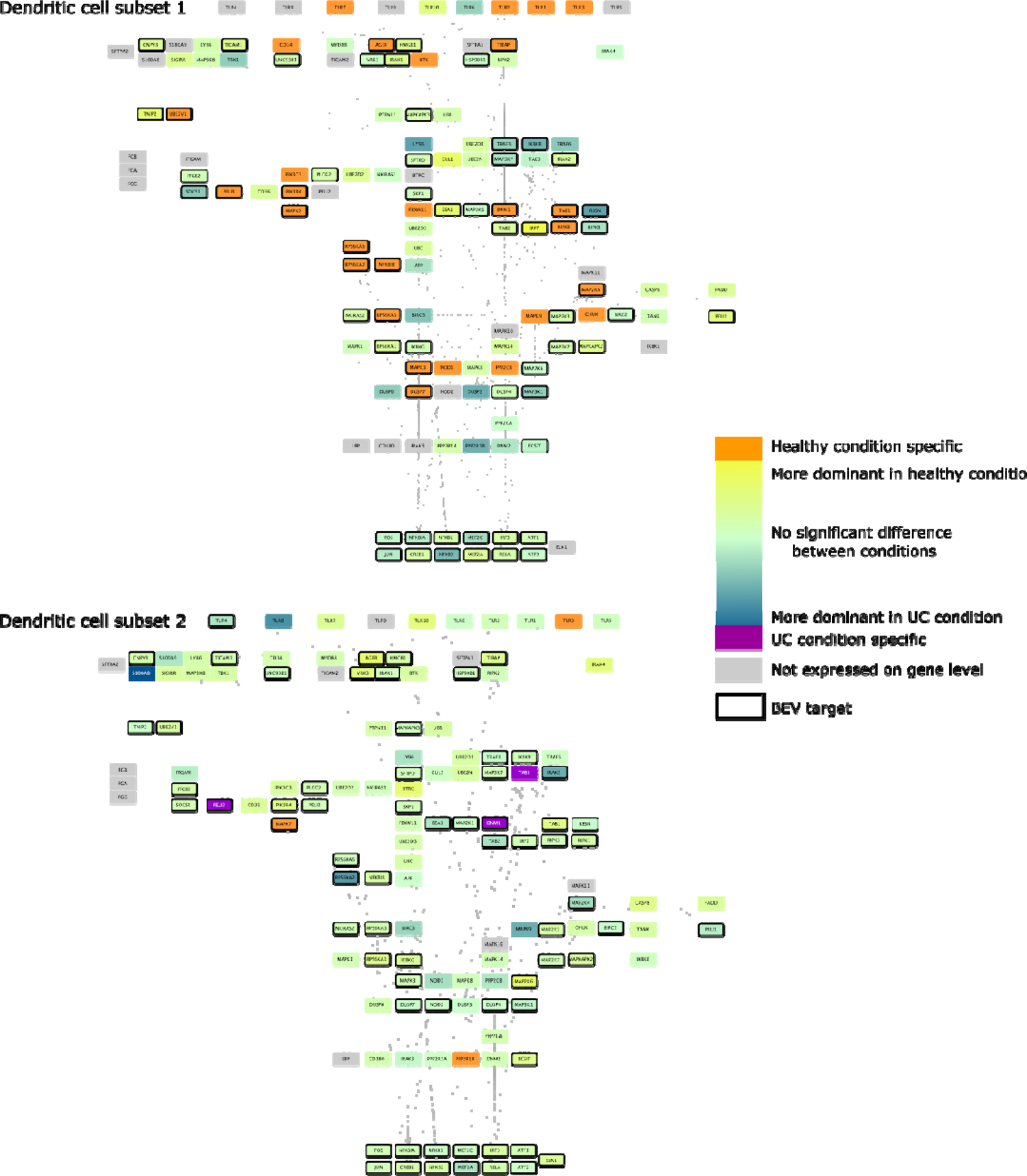
TLR pathway in DCs. Edges between nodes represent protein-protein interactions. Figures have been created with Cytoscape [32]

Analysis of TLR pathways demonstrated cell specificity, especially in monocytes and DC1 cells, with differences occurring mostly at the level of receptor proteins and in terms of transcription factors (TFs), gene expression did not show divergence between the healthy and diseased conditions. Based on these results, proteins in BEVs are more connected to downstream proteins and TFs than to receptors.

Dendritic cell subsets (DC1-DC2) show diverse characteristics regarding expression of TLR pathway members. In DC1 cells under healthy conditions, a large number of TLR pathway members were expressed in a condition-specific manner, including TLR1, 2, 3 and 7. In DC2 cells, three-three proteins were uniquely found in healthy (TLR3, MAPK7, and PP2R1B) and in non-inflamed UC (TAB3, DNM1, and PELI3) conditions. In addition, more TLR receptors (TLR1-8, TLR10) were represented in DC2 cells compared to DC1 cells. However, no significant differences were detected in the expression of TLR pathway members between the different conditions, unlike in DC1 cells. While no receptor was targeted in DC1, TLR4 was identified as a potential BEV target in DC2 cells [Figure 4].

In monocytes, the majority of TLR pathway members were expressed with signals being spread through diverse paths due to a few key signalling proteins being represented only in the healthy or diseased network. In terms of cycling monocytes, TLR1,2,5,6,7,8 were expressed at equivalent levels in both conditions, with TLR4 strongly related to the diseased condition.

Amongst downstream signalling components, nine proteins were represented in the healthy state and two proteins in non-inflamed UC with BEV proteins being able to bind almost all of them. In inflammatory monocytes several condition-specific pathways were identified including TLR4 and TLR5 in non-inflamed UC, and TLR7 and TLR10 pathways in the healthy state. The network shows a high number of condition-specific proteins downstream (17 healthy and 12 UC specific proteins) [Figure 5]. These results show that BEV proteins bind one TLR receptor (TLR4) which is expressed in both cell-types but only in inflammation-related monocytes in non-inflamed UC.

**Figure 5:**
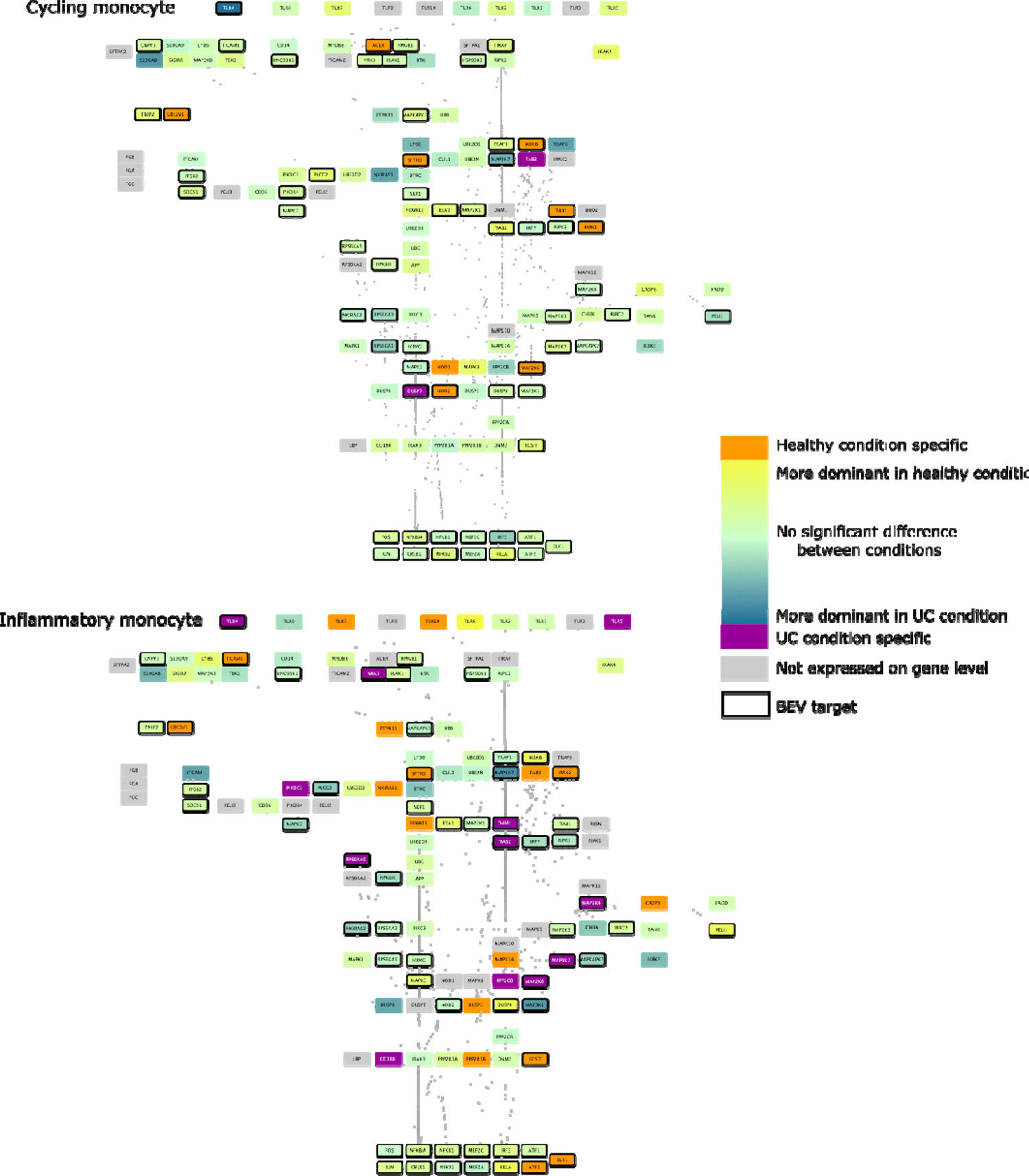
TLR pathway in monocytes. Edges between nodes represent protein-protein interactions. Figures have been created with Cytoscape [32].

We analysed bulk RNAseq datasets to verify the role of BEVs on the TLR4 pathway in THP-1 cell line (THP-1 monocyte cell line derived from human leukemic line [33]). Results showed a more similar network to the output of the cycling monocyte scRNAseq data analysis. However, we found a few differences in TLR pathways, revealing more potential for BEV-interacting proteins (PELI2-3, IRAK2, DNM1, RPS6K2, MAPK11) [Figure 6].

**Figure 6:**
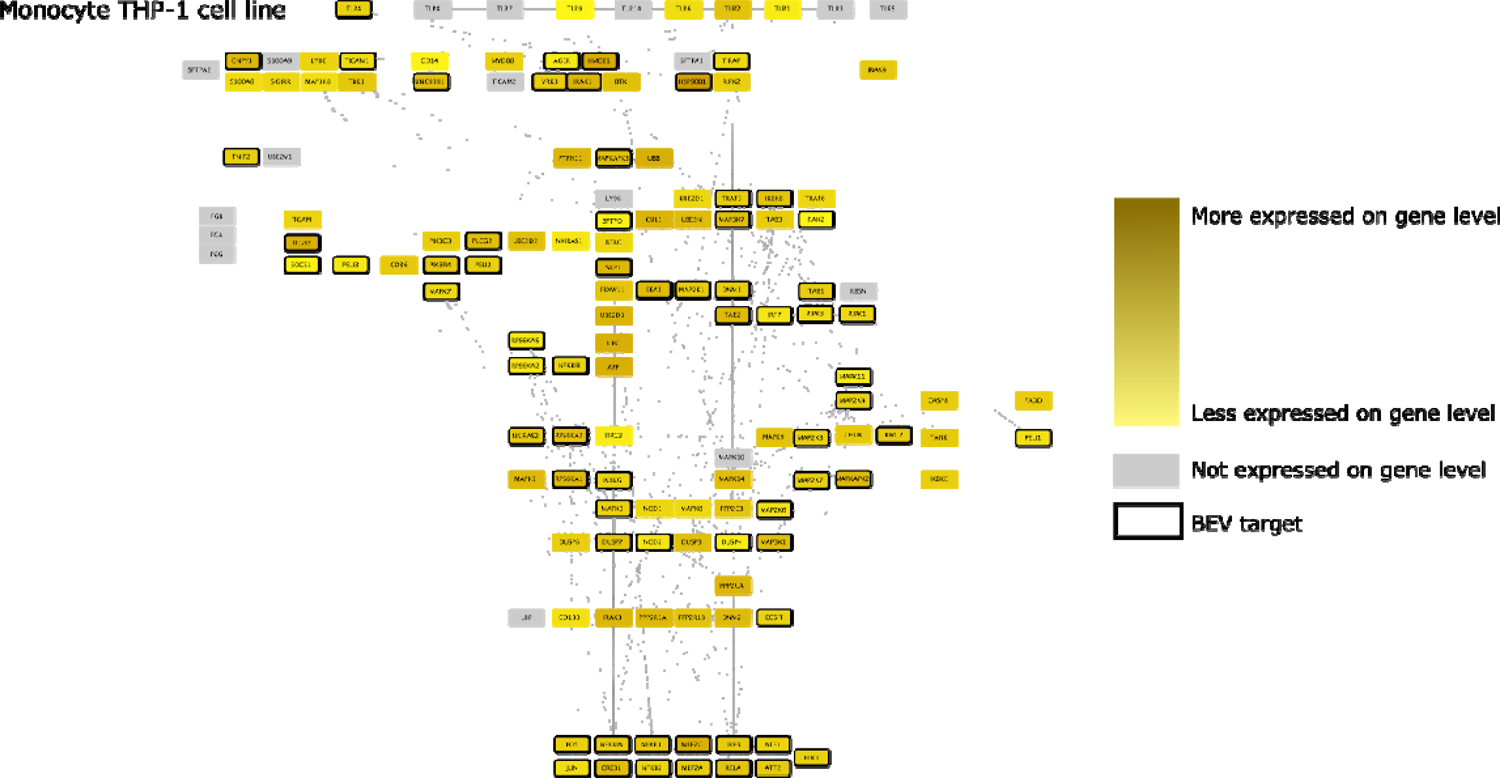
TLR pathway in THP-1 monocytes (based on bulk transcriptomic datasets). Edges between nodes represent protein-protein interactions. Figures have been created with Cytoscape [32].

Based on the pipeline, macrophages depict no significant alteration in UC compared to the healthy state. While 9/10 receptors are potentially represented, TLR4 was the only candidate interacting with BEV proteins. MAPK10-11 helped spread the signal in healthy cells, while PELI2 was expressed only in diseased macrophages [Figure 7].

**Figure 7:**
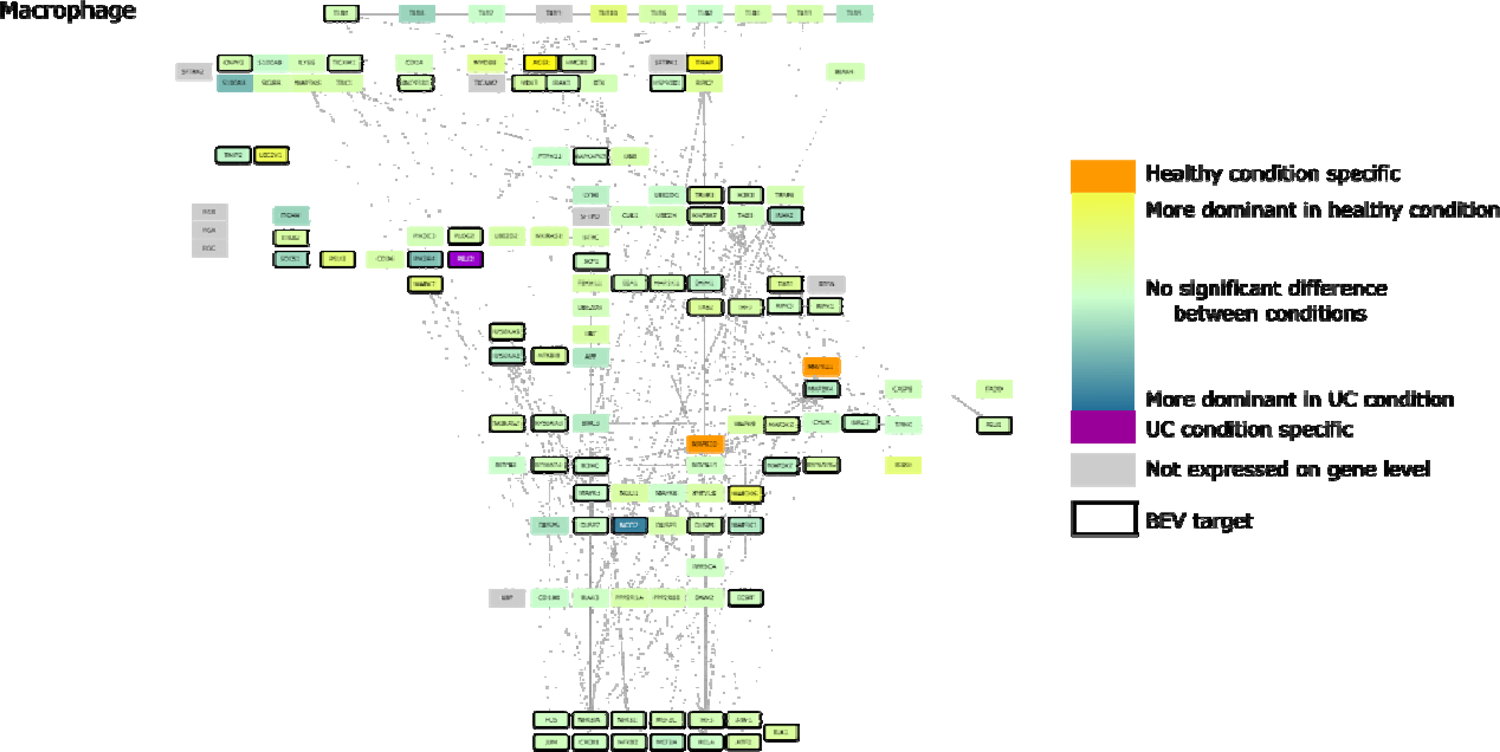
TLR pathway in macrophages. Edges between nodes represent protein-protein interactions. Figures have been created with Cytoscape [32].

### Inhibition of TLR4 signalling abrogates BEV-driven monocyte activation

Our pipeline identified TLR4 as the only receptor associated with BEVs in cycling monocytes, DC1 and macrophage cells, therefore we investigated the effects of BEVs on TLR-mediated activation of monocytes through BEV-monocyte co-culture experiments. Specifically, we focused on the role of TLR4 in the interaction between BEVs and monocytes/macrophages by co-culturing THP1 cells expressing an NF-kB reporter gene (THP1-Blue) with Bt BEVs. THP1-Blue cells were incubated with serial dilutions of Bt BEVs (3×10^9^ to 3×10^6^/mL) in the presence or absence of the TLR4 inhibitor, CLI-095 [34, 35], which in pre-optimisation experiments was shown to selectively inhibit TLR4 mediated activation of NF-kB (data not shown). THP1-Blue cells exposed to CLI-095 prior to incubation with 3×10^8^ and 3×10^7^ BEVs/mL demonstrated significant reduction of NF-kB activation (33.7% and 36.7% inhibition, respectively). The inhibitory effect of CLI-095 was lessened by excess of BEVs, as well as by low BEV concentrations insufficient to induce significant NF-kB activation [Figure 8]. These experimental findings validate our *in silico* analysis, identifying TLR4 as a BEV-interacting receptor. However, the inability to completely inhibit BEV-induced THP-1 activation by CLI-095 indicates a TLR4-independent effect of BEVs on NF-kB-activating pathways. This potential impact is shown on the TLR signalling network by highlighting the BEV interacting downstream pathway components.

**Figure 8:**
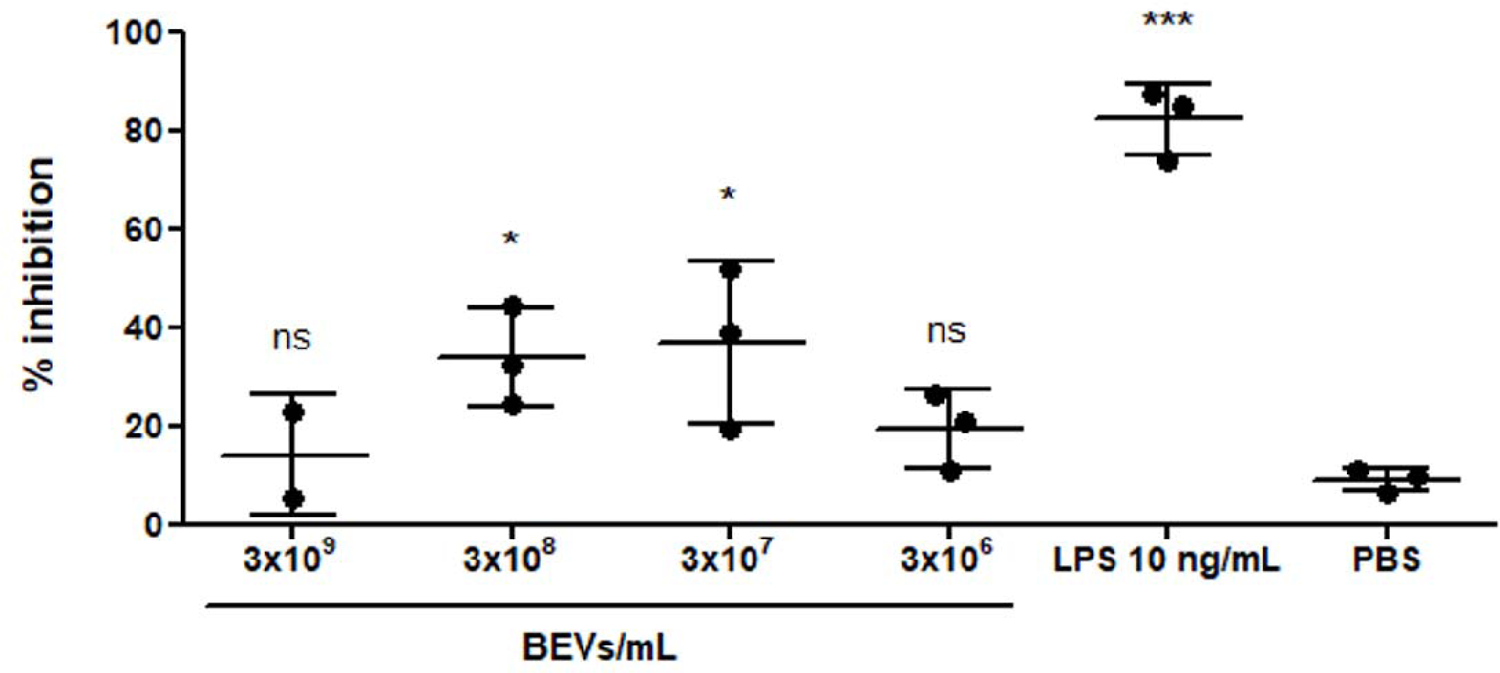
Inhibition of TLR4 signalling pathway abrogates THP1-Blue cells activation by Bt BEVs. NF-B activationκ promoted by different doses of BEVs in 5×10^5^ THP1-Blue cells/mL in the presence or absence of the TLR4 inhibitor CLI-095 was determined by measuring absorbance at 620 nm after incubation with the colorimetric assay reagent Quanti-Blue, using LPS from *E. coli* as positive control and PBS as negative control. Data are presented as percentage of inhibition calculated as the ratio between the absorbance at 620 nm of each sample incubated in the presence/absence of the TLR4 inhibitor (mean ± SD, n=3). Significant differences between each experimental group and the negative control were L determined by using one-way ANOVA followed by Dunnett’s multiple comparison post hoc test. * (p < 0.05), *** L L (p<0.001), **** (p < 0.0001).

## Discussion

BEVs contain proteins capable of affecting the host immune system [36]. However, the molecular mechanisms of BEV-mediated signalling in host cells are poorly understood. The recent availability of scRNAseq data facilitates the analysis of biological pathways at cell-type specific resolution, which we utilised here to identify the differential effects of BEV exposure on host immune cells.

Specifically, we examined proteins in BEVs generated by the major human commensal gut bacterium, Bt, which is a potential therapeutic agent in IBD [1]. Hence, it is important to understand which cell-types are affected by Bt BEVs. Considering gene expression profiles are different not only among cells but also in the same cells under different conditions, the possible protein-protein interactions will vary between microbes and its host.

To investigate the BEV-host cell interactome we developed a novel computational pipeline combining single-cell transcriptomics with network biology approaches to reconstruct the interactomes and model the effect of Bt BEVs on different types of immune cells. We used publicly available human scRNAseq data to examine how Bt BEVs could potentially impact cycling monocytes, inflammatory monocytes, DC1s, DC2s, and macrophages in both the healthy, disease-free, colon and non-inflamed UC colon [19]. The output of the workflow highlighted that Bt BEVs have a large number of interactions with these immune cells. The majority of candidate interacting BEV proteins are catabolic enzymes with numerous non-specific connections. We did however identify microbial helicases targeting the human polymerase protein PAPD5. Binding of helicases to polymerase proteins is critical to initiate leading-strand DNA synthesis [37]. PAPD5 is also a well-known negative regulator of miR-21. Among the targets of this miRNA are genes involved in the immune responses and pathogenesis of autoimmune diseases, including IBD [38, 39].

Despite the large overlap of connections, we identified among five immune cell-types unique functions triggered by Bt BEVs in the healthy and UC colon. For example, cell division is significantly enriched in cycling monocytes in the healthy state. In healthy conditions, bone marrow-derived monocytes circulate in the blood and differentiate to macrophages in various tissues. Therefore, the proliferation of monocytes is required to maintain a pool of tissue specific monocytes/macrophages [40]. We also inferred that in UC DNA repair activity might be influenced by BEV proteins interacting with cycling monocytes. Prior work has demonstrated that patients with UC have higher levels of mucosal oxidative DNA damage, even under non-inflamed conditions, which increases with the duration and severity of disease [41–45]. This is a potential explanation for the higher incidence of colorectal cancer in UC patients. Indeed, mice with chronic colitis that are deficient in a key DNA repair enzyme have increased susceptibility to developing colorectal carcinoma in response to oxidative stress [46]. Our findings suggest Bt BEV proteins may play an important role in promoting DNA repair activity against oxidative DNA damage in cycling monocytes in patients with UC.

In inflammatory monocytes, BEV proteins upregulate apoptosis and the ERAD pathway in both healthy and UC states. Both these cellular processes are critical components of the unfolded protein response (UPR), which is important for resolving ER stress. Interestingly, our analysis also showed that BEVs influence ER-stress response related processes in most of the immune cell-types we studied. In UC, several risk variants affect genes involved in these pathways and together with environmental factors (such as dysbiosis, metabolites and/or inflammatory cytokines), disrupt the UPR in intestinal epithelial cells. The resultant unabated ER stress has been shown to precipitate intestinal inflammation. However, in monocytes and macrophages higher levels of UPR transcripts have been found in DSS-colitis mice compared to control mice, suggesting that the UPR may permit these cells to survive in the inflamed mucosal milieu of colitis [47]. Thus, BEV proteins may help promote resolution of ER stress and maintain the survival of inflammatory monocytes, macrophages, and other immune cells by upregulating key components of the UPR.

Dendritic cells are important antigen presenting cells and play important roles in innate and adaptive immunity including responses to microbial pathogens. Interactions between DCs and BEVs can direct inflammation in the gut [12]. The microbiome composition can lead to the differentiation of immature DCs into diverse subpopulations therefore maintaining immune homeostasis [48] We next focused on the effects of BEVs on TLR pathways examining different activities and outcomes of the pathways. Literature evidence indicates that Bt is capable of binding the TLR4 receptor [49]. While the study examined the mouse intestine, here we discovered the same interaction on protein level in a cell-type and condition specific manner in human. Regarding the role of TLR4 in UC, the receptor is upregulated in the inflamed colonic mucosa of UC patients identified by increased mRNA and protein levels [50, 51]. The TLR4 receptor transmits signals via MYD88-dependent and MYD88-independent pathways. In the MYD88-dependent pathway, MYD88 and the IRAK complex activate TAK1 and IKK. Consequently, NF-βB translocates to the nucleus, and with FOS and JUN transcription factors enhances the κ expression of pro-inflammatory cytokines [52].

Based on our Bt BEV-human PPI network, a bacterial carboxyl-terminal protease is predicted to bind the TLR4 receptor. There is however no evidence as to how this enzyme affects TLR4 receptor activation, although in chickens the TLR15 receptor can be triggered by microbial proteases [53]. The BEV-host interactome analysis also highlighted IRAK1-4, IKBKB, NFKB1-2 as potential BEV interacting host proteins, suggesting Bt may bind not only the receptor itself but also downstream intracellular components of TLR4 pathway. The co-localization of Bt BEVs with various intracellular compartments and in particular, the nucleus, in intestinal epithelial cells that have acquired BEVs [7] demonstrates the feasibility of Bt BEVs interactions with various cytoplasmic constituents of host cells.

Our study highlighted the importance of both condition and cell specific analyses. The signalling in inflammatory monocytes showed that TLR4 is expressed only in the diseased condition. In healthy, non-inflamed, conditions this subset of monocytes represents a small leukocyte subset in the blood, although they are dominant during inflammation [54]. Regarding cell specificity, TLR4 expressed in inflammation-related subpopulation of DCs and not in healthy conditions, highlighting the difference between immune cell subsets. The pipeline emphasizes the importance of using scRNAseq data to understand the molecular basis of BEV protein effects on host immune cells.

We confirmed the role of BEVs in TLR4-mediated activation of human monocytes using the THP-1 human monocytic cell line using bulk transcriptomic datasets for the cell line. Although the bulk RNAseq data identified less expressed TLR receptors but more BEV interacting downstream components, it was broadly consistent our findings based on scRNAseq data. Bt BEVs activate the TLR4 pathway, as shown using the TLR4-specific inhibitor CLI-095, which reduced the BEV-induced NF-kB activation. The inability to completely inhibit BEV-induced NF-kB activation indicates that other PRR NF-kB-activating receptors are involved in BEV-monocyte interactions. This is consistent with our results from the bioinformatic analysis, where proteins from BEVs can bind to several downstream members of the TLR intracellular signalling pathway.

Whilst providing new and potentially important insights into BEV-host immune cell interactomes our pipeline is limited to including only one available scRNAseq dataset that describes gene expression in healthy and non-inflamed UC colonic mucosal cells. Some expressed genes could be missed with the 10X single-cell transcriptomics approach, and we do not have corresponding protein levels (or their activities) in the cells of interest. In inferring a microbe-host PPI network, we assumed that all genes were translated to functional proteins regardless of post-transcriptional modifications that could affect protein abundance. Regarding the PPI predictions for the microbe-host interactions we used a limited list of domain-motif interactions from the ELM database. Finally, our workflow cannot predict the activation or inhibitory effects of BEV proteins, but only whether they act on a particular receptor and pathway. Further investigations are needed to establish the binding mechanism and impact of for example, the BEV carboxy-peptidase on host TLR4 receptors. Despite these limitations, our pipeline provides a deeper insight into the effect of BEV proteins on host immunity on protein level and shows the importance of condition and cell specificity. We also not only predicted the affected host processes that are supported by the literature but also identified new potential targets for experimental validation.

## Conclusion

We have developed a computational pipeline that predicts both the cell and condition specific effects of Bt BEV proteins on key host immune cell populations. Focusing on the inflammation-related TLR pathway, which plays a role in IBD pathogenesis, our workflow highlighted the importance of single-cell based analysis identifying differences in TLR4 receptor expression in diverse DC subpopulations. The current pipeline offers potentially interesting connection points between Bt and host immune cells that will aid in understanding how BEVs and their protein cargo may be of therapeutic value in IBD.

## Acknowledgement

This work was supported by the UKRI Biotechnological and Biosciences Research Council (BBSRC) UK grant awarded to the Earlham Institute (BB/J004529/1, BB/P016774/1, and BB/CSP17270/1) and to the Quadram Institute’s Gut Microbes and Health Institute Strategic Programme (BB/R012490/1, BBS/E/F/000PR10353 and BBS/E/F/000PR10355). LG was supported by the BBSRC Norwich Research Park Biosciences Doctoral Training Partnership grant number BB/M011216/1. JPT is an Academic Clinical Fellow funded by the National Institute of Health Research (NIHR).

## Supplementary data

**Supplementary Figure 1:**
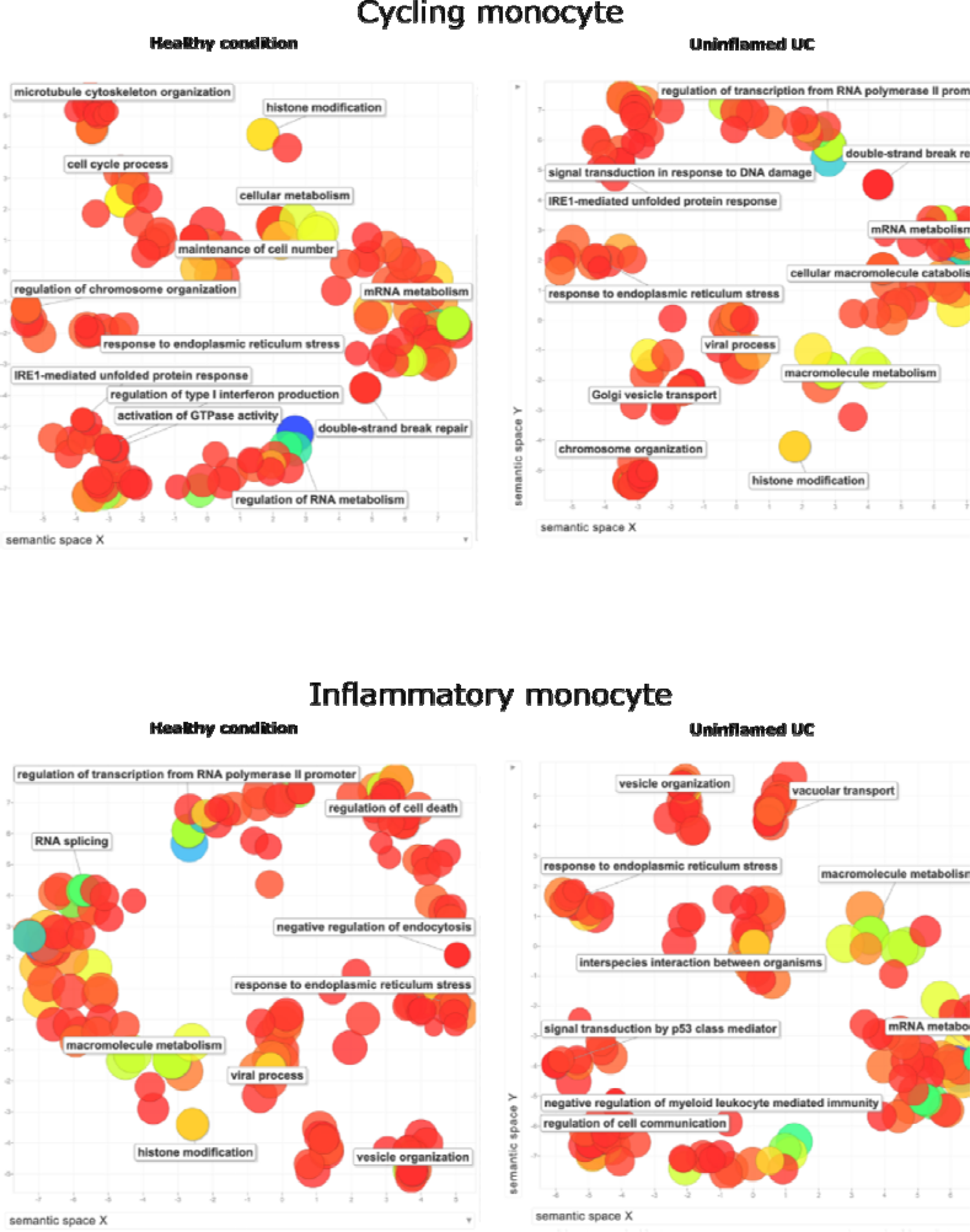
Functional analysis of BEV targets in monocytes. Size of the points is equal to the number of proteins involved in the function, the colour represents the log10 p-value (blue - lowest value, red - highest value)

**Supplementary Figure 2:**
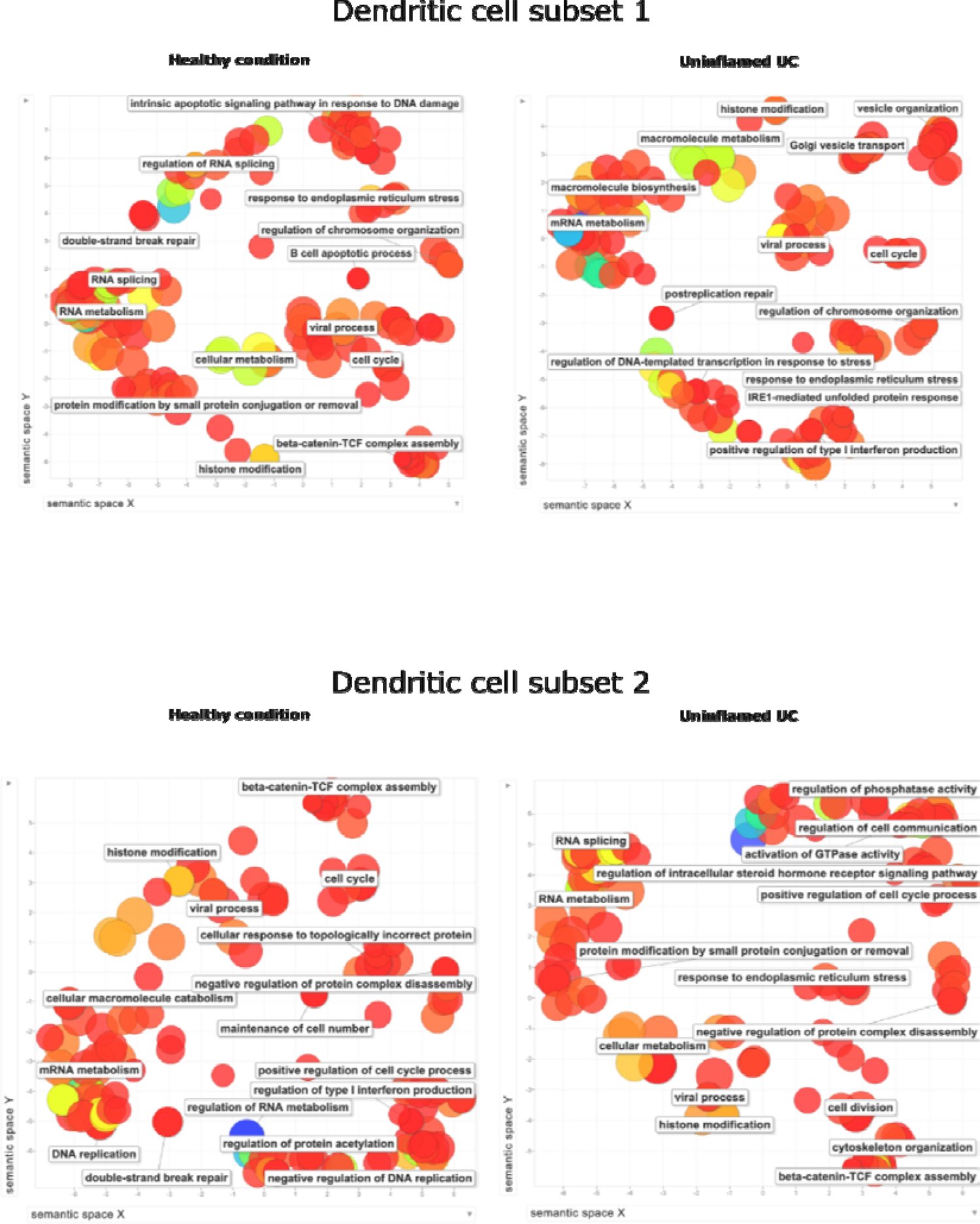
Functional analysis of BEV targets in dendritic cells. Size of the points is equal to the number of proteins involved in the function, the colour represents the log10 p-value (blue - lowest value, red highest value)

**Supplementary Figure 3:**
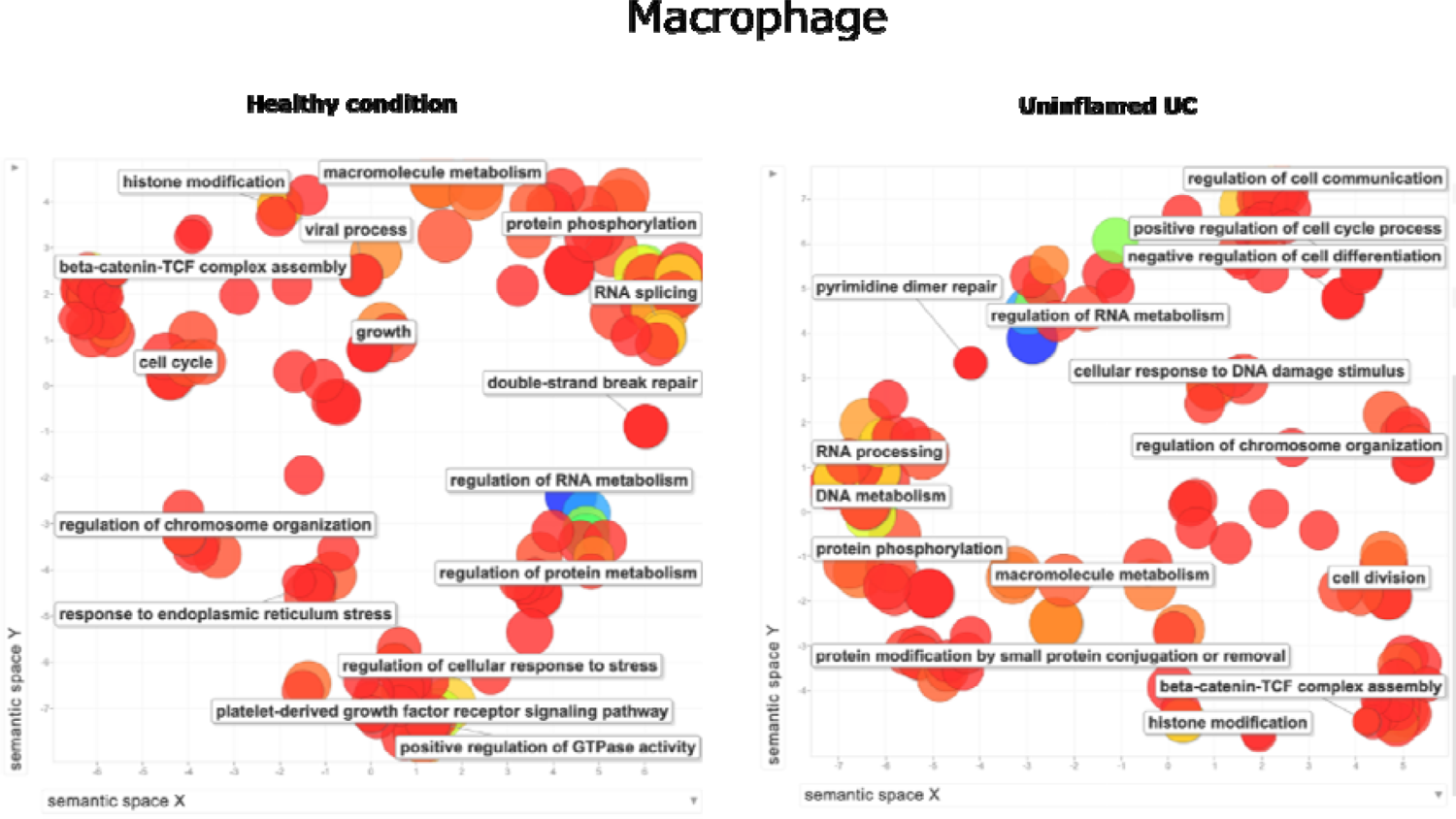
Functional analysis of BEV targets in macrophages. Size of the points is equal to the number of proteins involved in the function, the colour represents the log10 p-value (blue - lowest value, red highest value)

**Supplementary Table 1.**
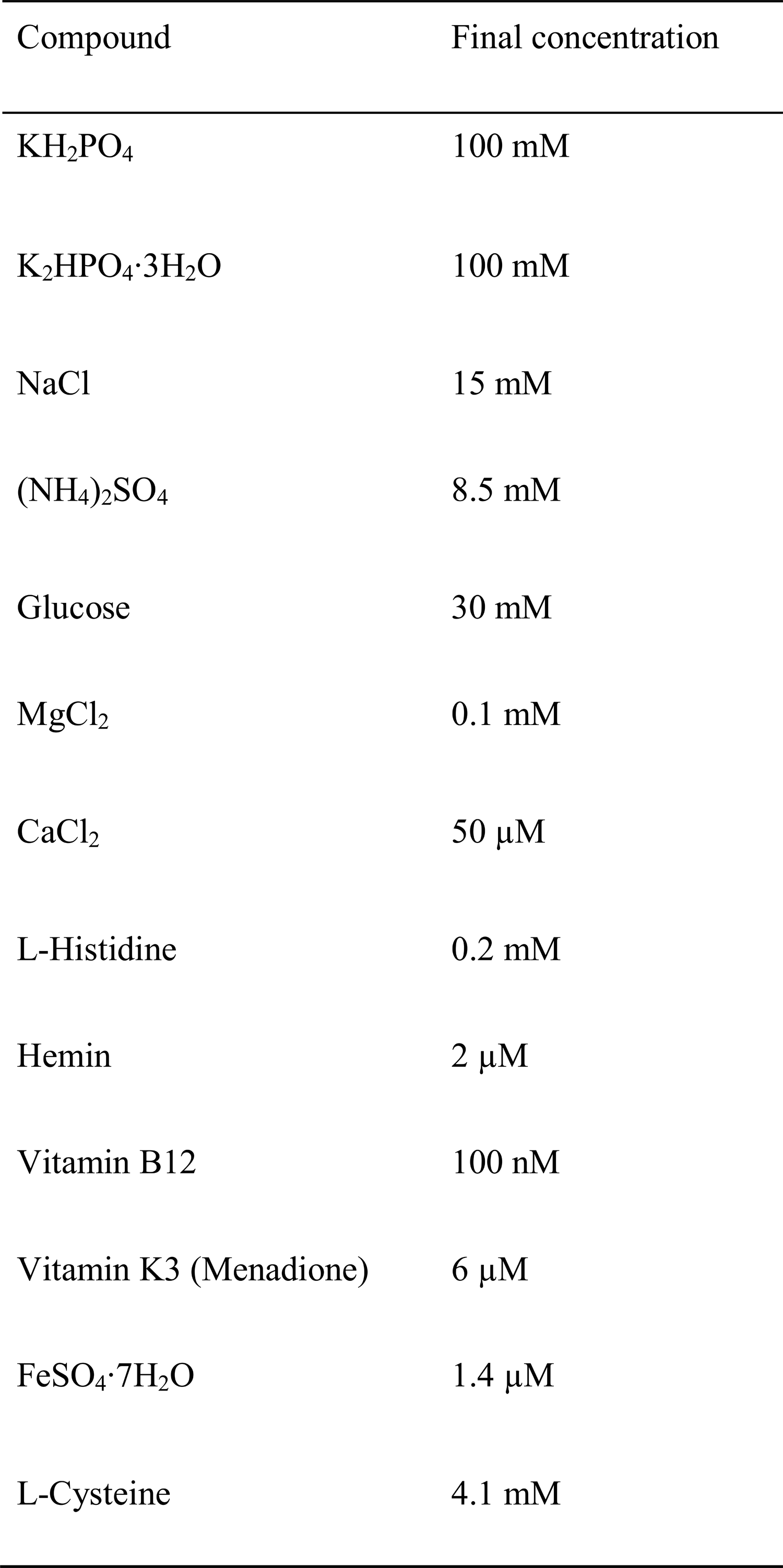
BDM+ composition

**Supplementary Table 2.**
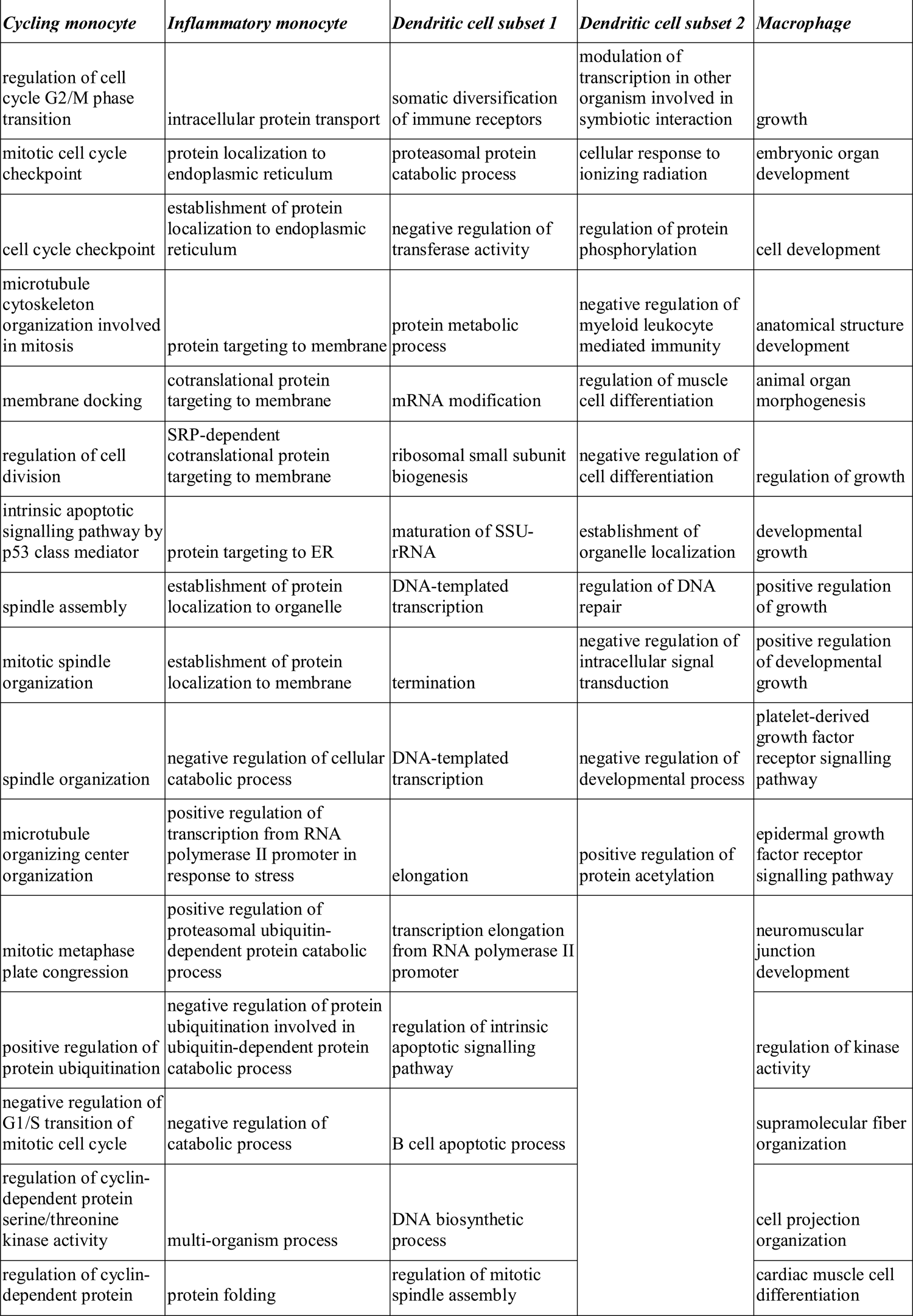
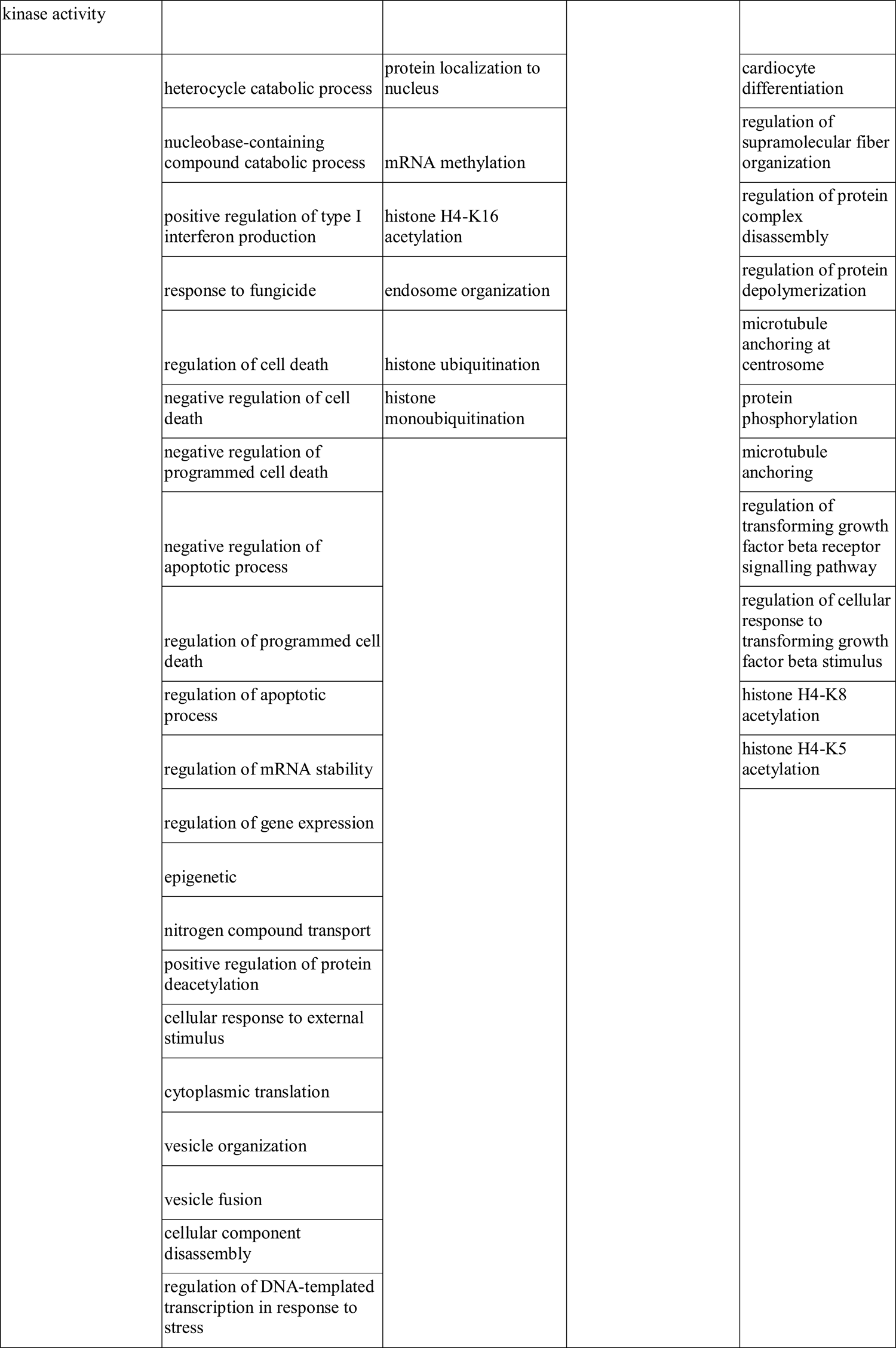
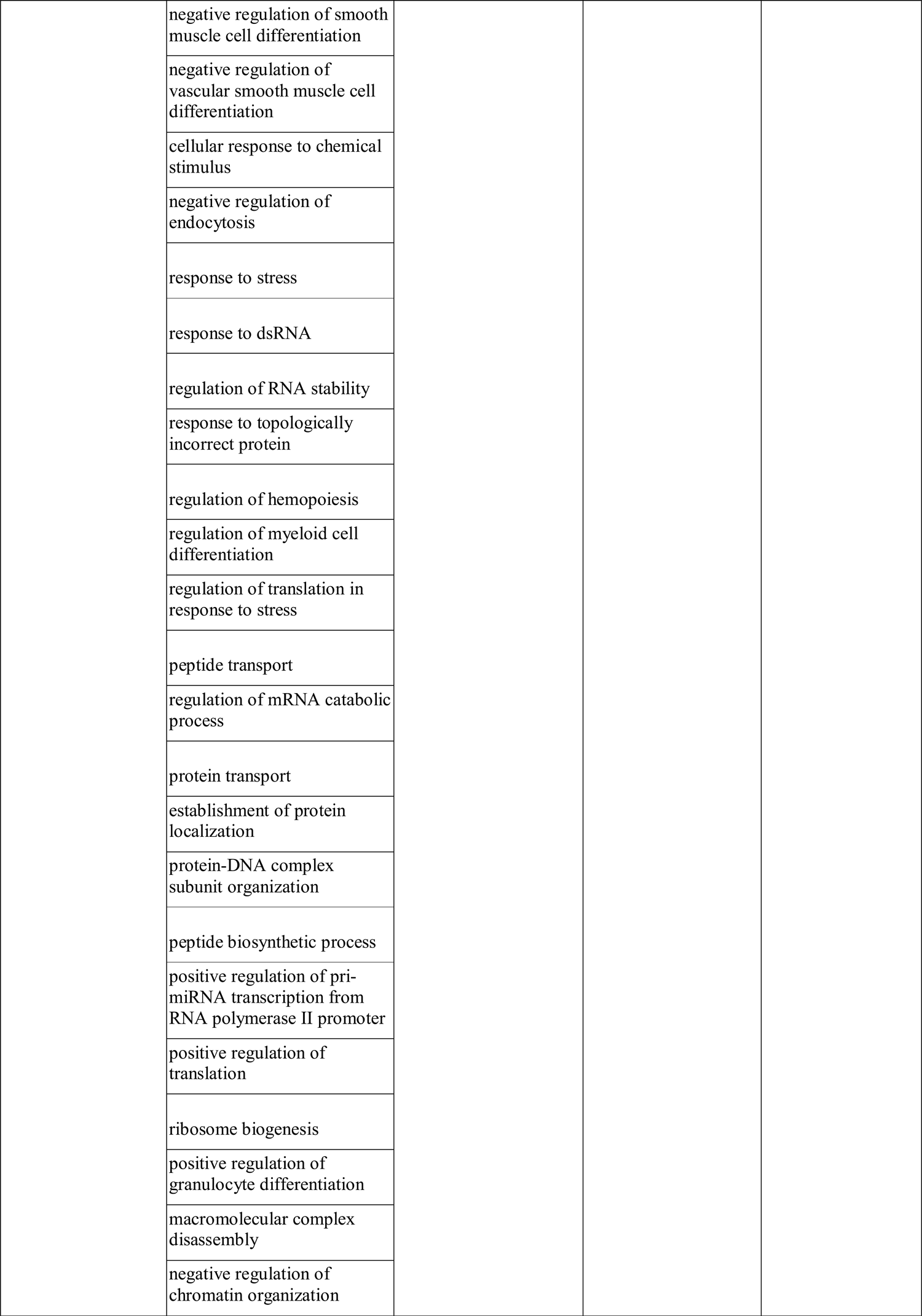
Unique annotations in cells affected by BEV proteins in healthy condition

**Supplementary Table 3.**
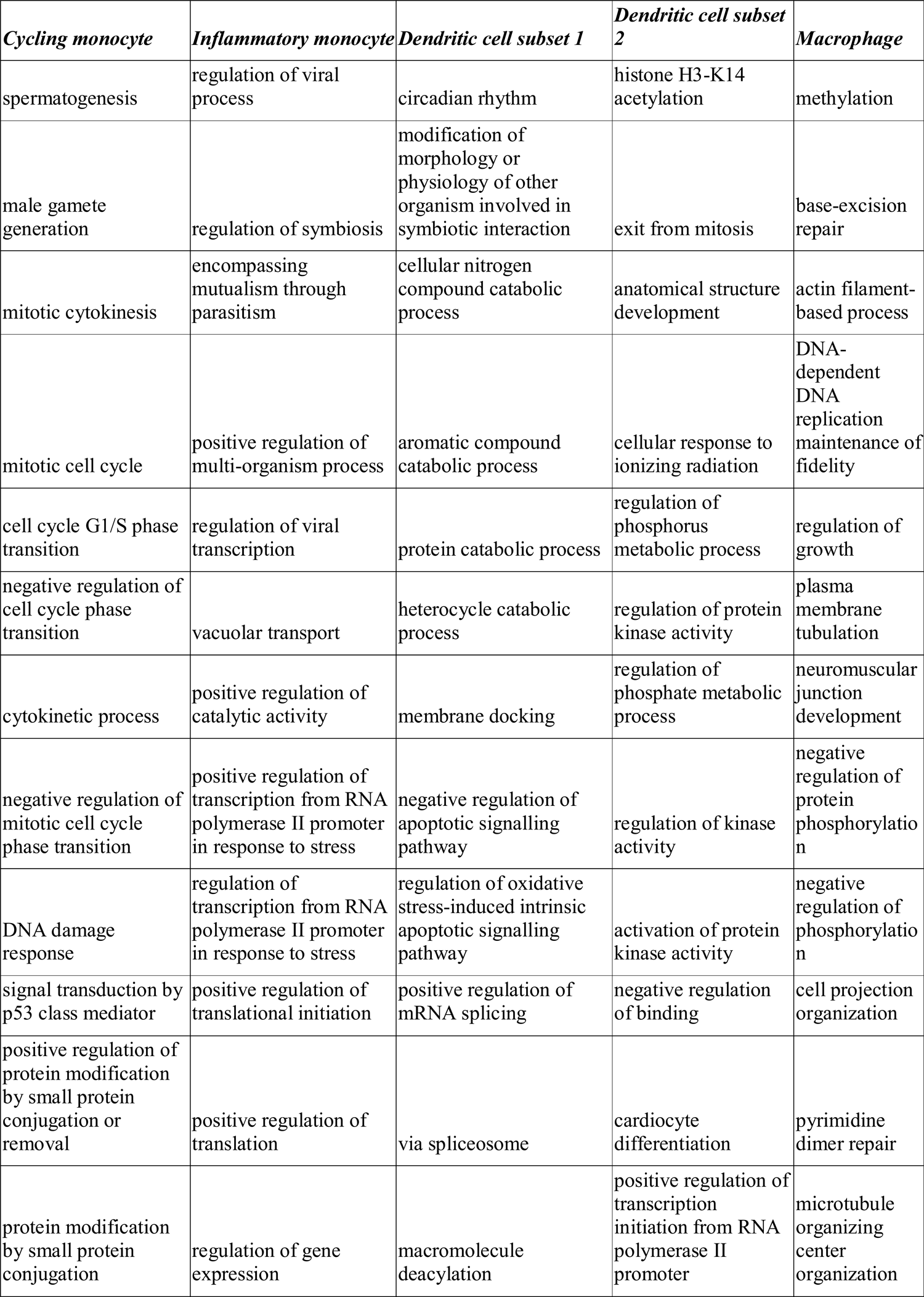
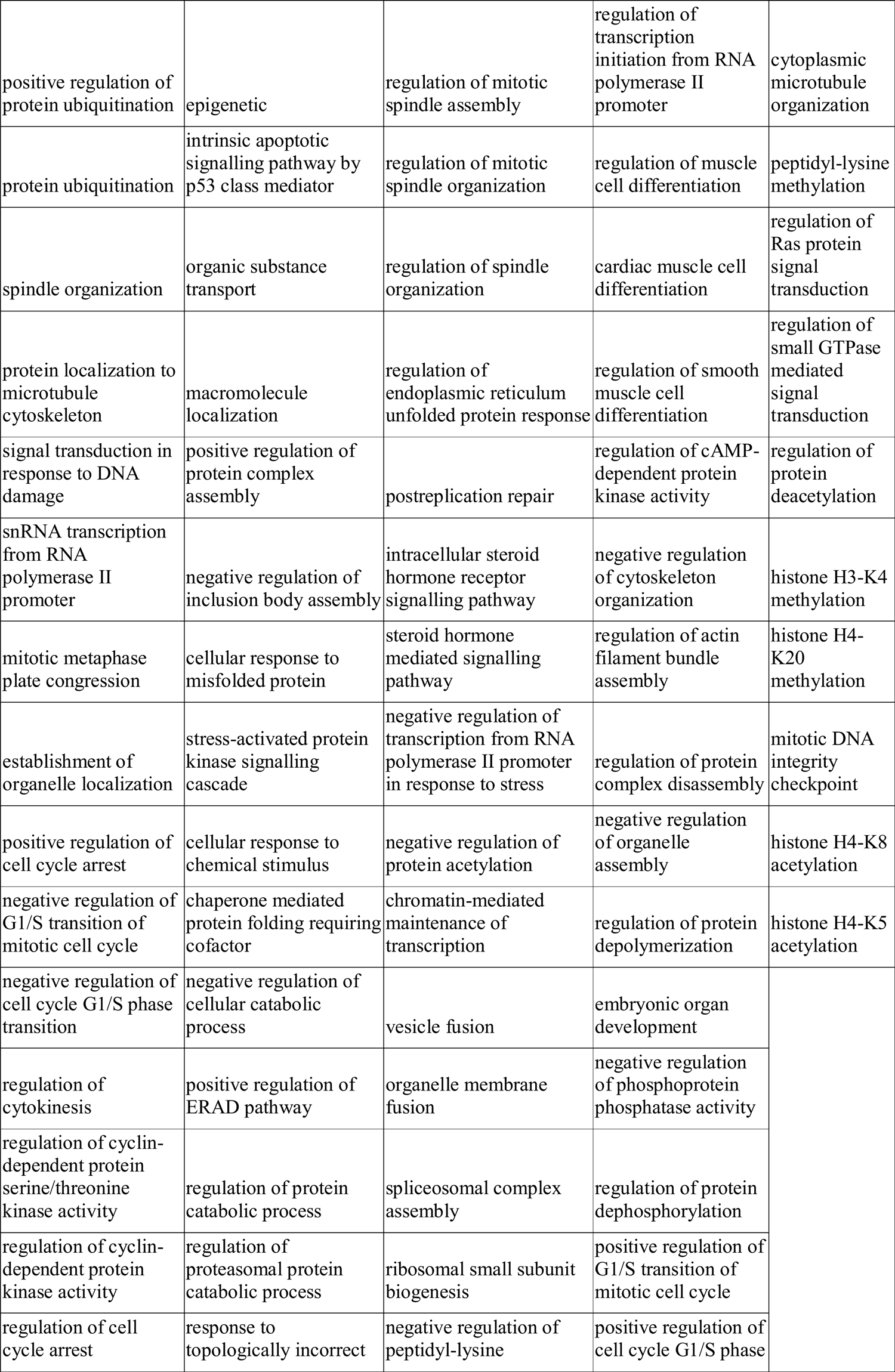
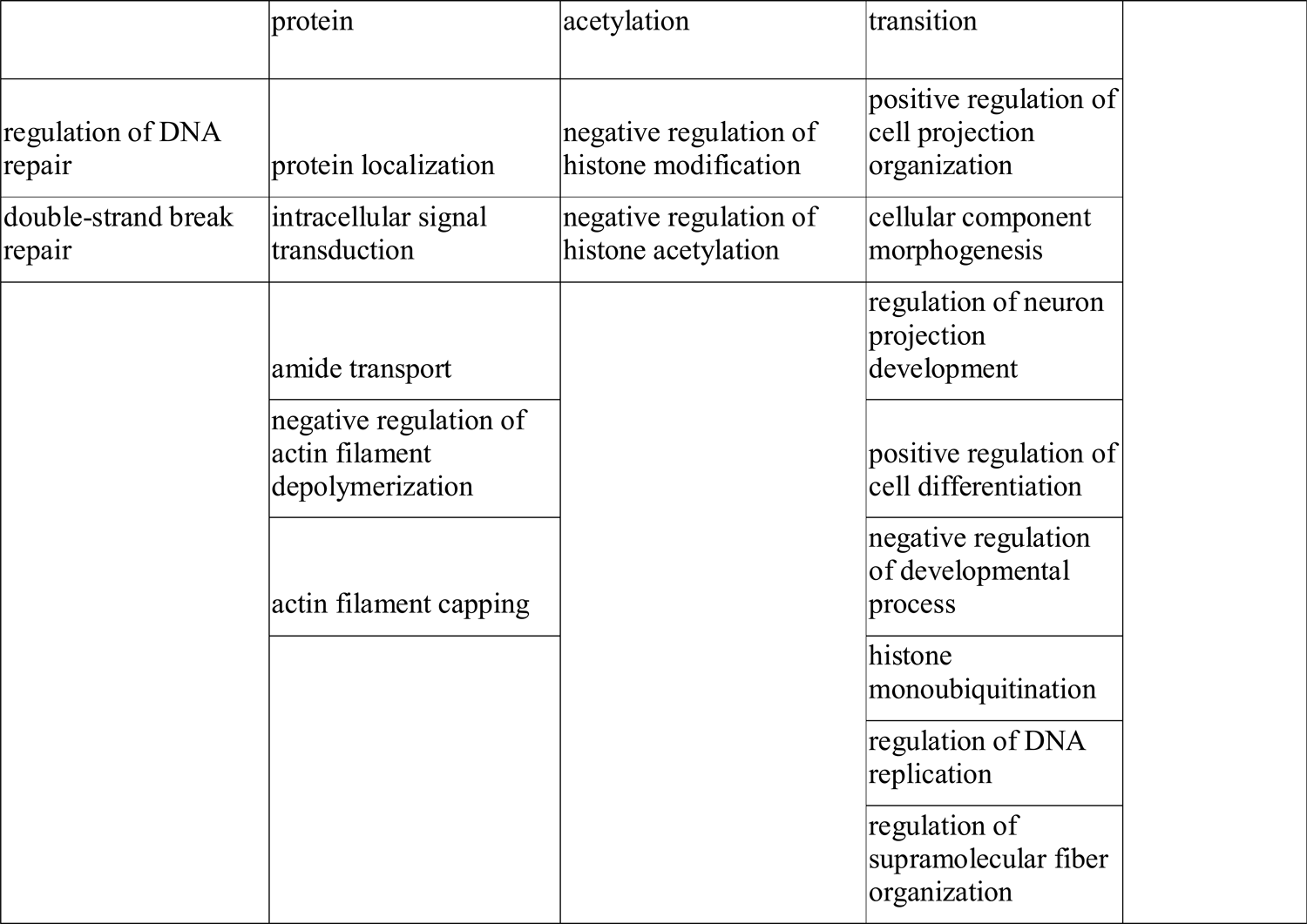
Unique annotations in cells affected by BEV proteins in UC condition

## References

1. Delday M, Mulder I, Logan ET, Grant G. Bacteroides thetaiotaomicron Ameliorates Colon Inflammation in Preclinical Models of Crohn’s Disease. Inflamm Bowel Dis. 2019;25:85–96. doi:10.1093/ibd/izy281.

2. Kabeerdoss J, Jayakanthan P, Pugazhendhi S, Ramakrishna BS. Alterations of mucosal microbiota in the colon of patients with inflammatory bowel disease revealed by real time polymerase chain reaction amplification of 16S ribosomal ribonucleic acid. Indian J Med Res. 2015;142:23–32. doi:10.4103/0971-5916.162091.

3. Bacteroides Thetaiotaomicron - an overview | ScienceDirect Topics. https://www.sciencedirect.com/topics/immunology-and-microbiology/bacteroides-thetaiotaomicron. Accessed 23 Feb 2021.

4. Chang X, Wang S-L, Zhao S-B, Shi Y-H, Pan P, Gu L, et al. Extracellular Vesicles with Possible Roles in Gut Intestinal Tract Homeostasis and IBD. Mediators Inflamm. 2020;2020:1945832. doi:10.1155/2020/1945832.

5. Fábrega M-J, Rodríguez-Nogales A, Garrido-Mesa J, Algieri F, Badía J, Giménez R, et al. Intestinal Anti-inflammatory Effects of Outer Membrane Vesicles from Escherichia coli Nissle 1917 in DSS-Experimental Colitis in Mice. Front Microbiol. 2017;8:1274. doi:10.3389/fmicb.2017.01274.

6. Schwechheimer C, Kuehn MJ. Outer-membrane vesicles from Gram-negative bacteria: biogenesis and functions. Nat Rev Microbiol. 2015;13:605–19. doi:10.1038/nrmicro3525.

7. Jones EJ, Booth C, Fonseca S, Parker A, Cross K, Miquel-Clopés A, et al. The Uptake, Trafficking, and Biodistribution of Bacteroides thetaiotaomicron Generated Outer Membrane Vesicles. Front Microbiol. 2020;11:57. doi:10.3389/fmicb.2020.00057.

8. Shen Y, Giardino Torchia ML, Lawson GW, Karp CL, Ashwell JD, Mazmanian SK. Outer membrane vesicles of a human commensal mediate immune regulation and disease protection. Cell Host Microbe. 2012;12:509–20. doi:10.1016/j.chom.2012.08.004.

9. Hickey CA, Kuhn KA, Donermeyer DL, Porter NT, Jin C, Cameron EA, et al. Colitogenic Bacteroides thetaiotaomicron Antigens Access Host Immune Cells in a Sulfatase-Dependent Manner via Outer Membrane Vesicles. Cell Host Microbe. 2015;17:672–80. doi:10.1016/j.chom.2015.04.002.

10. Kaparakis-Liaskos M, Ferrero RL. Immune modulation by bacterial outer membrane vesicles. Nat Rev Immunol. 2015;15:375–87. doi:10.1038/nri3837.

11. Cecil JD, Sirisaengtaksin N, O’Brien-Simpson NM, Krachler AM. Outer Membrane Vesicle-Host Cell Interactions. Microbiol Spectr. 2019;7. doi:10.1128/microbiolspec.PSIB-0001-2018.

12. Durant L, Stentz R, Noble A, Brooks J, Gicheva N, Reddi D, et al. Bacteroides thetaiotaomicron-derived outer membrane vesicles promote regulatory dendritic cell responses in health but not in inflammatory bowel disease. Microbiome. 2020;8:88. doi:10.1186/s40168-020-00868-z.

13. Stentz R, Carvalho AL, Jones EJ, Carding SR. Fantastic voyage: the journey of intestinal microbiota-derived microvesicles through the body. Biochem Soc Trans. 2018;46:1021–7. doi:10.1042/BST20180114.

14. Kawasaki T, Kawai T. Toll-like receptor signaling pathways. Front Immunol. 2014;5:461. doi:10.3389/fimmu.2014.00461.

15. Zhang M, Sun K, Wu Y, Yang Y, Tso P, Wu Z. Interactions between Intestinal Microbiota and Host Immune Response in Inflammatory Bowel Disease. Front Immunol. 2017;8:942. doi:10.3389/fimmu.2017.00942.

16. Scott NA, Mann ER. Regulation of mononuclear phagocyte function by the microbiota at mucosal sites. Immunology. 2020;159:26–38. doi:10.1111/imm.13155.

17. Steinbach EC, Plevy SE. The role of macrophages and dendritic cells in the initiation of inflammation in IBD. Inflamm Bowel Dis. 2014;20:166–75. doi:10.1097/MIB.0b013e3182a69dca.

18. Stentz R, Wegmann U, Guirro M, Bryant W, Ranjit A, Goldson AJ, et al. Extracellular vesicles released by the human gut symbiont Bacteroides thetaiotaomicron in the mouse intestine are enriched in a selected range of proteins that influence host cell physiology and metabolism. 2020. doi:10.21203/rs.3.rs-124947/v1.

19. Smillie CS, Biton M, Ordovas-Montanes J, Sullivan KM, Burgin G, Graham DB, et al. Intra- and Inter-cellular Rewiring of the Human Colon during Ulcerative Colitis. Cell. 2019;178:714–730.e22. doi:10.1016/j.cell.2019.06.029.

20. Hart T, Komori HK, LaMere S, Podshivalova K, Salomon DR. Finding the active genes in deep RNA-seq gene expression studies. BMC Genomics. 2013;14:778. doi:10.1186/1471-2164-14-778.

21. Andrighetti T, Bohar B, Lemke N, Sudhakar P, Korcsmaros T. MicrobioLink: An Integrated Computational Pipeline to Infer Functional Effects of Microbiome-Host Interactions. Cells. 2020;9. doi:10.3390/cells9051278.

22. Kumar M, Gouw M, Michael S, Sámano-Sánchez H, Pancsa R, Glavina J, et al. ELM-the eukaryotic linear motif resource in 2020. Nucleic Acids Res. 2020;48:D296–306. doi:10.1093/nar/gkz1030.

23. UniProt Consortium. UniProt: a worldwide hub of protein knowledge. Nucleic Acids Res. 2019;47:D506–15. doi:10.1093/nar/gky1049.

24. Mészáros B, Erdos G, Dosztányi Z. IUPred2A: context-dependent prediction of protein disorder as a function of redox state and protein binding. Nucleic Acids Res. 2018;46:W329– 37. doi:10.1093/nar/gky384.

25. Eden E, Navon R, Steinfeld I, Lipson D, Yakhini Z. GOrilla: a tool for discovery and visualization of enriched GO terms in ranked gene lists. BMC Bioinformatics. 2009;10:48. doi:10.1186/1471-2105-10-48.

26. Supek F, Bošnjak M, Škunca N, Šmuc T. REVIGO summarizes and visualizes long lists of gene ontology terms. PLoS ONE. 2011;6:e21800. doi:10.1371/journal.pone.0021800.

27. Heberle H, Meirelles GV, da Silva FR, Telles GP, Minghim R. InteractiVenn: a web-based tool for the analysis of sets through Venn diagrams. BMC Bioinformatics. 2015;16:169. doi:10.1186/s12859-015-0611-3.

28. Jassal B, Matthews L, Viteri G, Gong C, Lorente P, Fabregat A, et al. The Reactome Pathway Knowledgebase. Nucleic Acids Res. 2020;48:D498–503. doi:10.1093/nar/gkz1031.

29. Türei D, Korcsmáros T, Saez-Rodriguez J. OmniPath: guidelines and gateway for literature-curated signaling pathway resources. Nat Methods. 2016;13:966–7. doi:10.1038/nmeth.4077.

30. Stentz R, Osborne S, Horn N, Li AWH, Hautefort I, Bongaerts R, et al. A bacterial homolog of a eukaryotic inositol phosphate signaling enzyme mediates cross-kingdom dialog in the mammalian gut. Cell Rep. 2014;6:646–56. doi:10.1016/j.celrep.2014.01.021.

31. Rammelt C, Bilen B, Zavolan M, Keller W. PAPD5, a noncanonical poly(A) polymerase with an unusual RNA-binding motif. RNA. 2011;17:1737–46. doi:10.1261/rna.2787011.

32. Shannon P, Markiel A, Ozier O, Baliga NS, Wang JT, Ramage D, et al. Cytoscape: a software environment for integrated models of biomolecular interaction networks. Genome Res. 2003;13:2498–504. doi:10.1101/gr.1239303.

33. Tsuchiya S, Yamabe M, Yamaguchi Y, Kobayashi Y, Konno T, Tada K. Establishment and characterization of a human acute monocytic leukemia cell line (THP-1). Int J Cancer. 1980;26:171–6. doi:10.1002/ijc.2910260208.

34. Ii M, Matsunaga N, Hazeki K, Nakamura K, Takashima K, Seya T, et al. A novel cyclohexene derivative, ethyl (6R)-6-[N-(2-Chloro-4-fluorophenyl)sulfamoyl]cyclohex-1-ene-1-carboxylate (TAK-242), selectively inhibits toll-like receptor 4-mediated cytokine production through suppression of intracellular signaling. Mol Pharmacol. 2006;69:1288–95. doi:10.1124/mol.105.019695.

35. Kawamoto T, Ii M, Kitazaki T, Iizawa Y, Kimura H. TAK-242 selectively suppresses Toll-like receptor 4-signaling mediated by the intracellular domain. Eur J Pharmacol. 2008;584:40–8. doi:10.1016/j.ejphar.2008.01.026.

36. Kuehn MJ, Kesty NC. Bacterial outer membrane vesicles and the host-pathogen interaction. Genes Dev. 2005;19:2645–55. doi:10.1101/gad.1299905.

37. Zhang H, Lee S-J, Zhu B, Tran NQ, Tabor S, Richardson CC. Helicase-DNA polymerase interaction is critical to initiate leading-strand DNA synthesis. Proc Natl Acad Sci USA. 2011;108:9372–7. doi:10.1073/pnas.1106678108.

38. Boele J, Persson H, Shin JW, Ishizu Y, Newie IS, Søkilde R, et al. PAPD5-mediated 3’ adenylation and subsequent degradation of miR-21 is disrupted in proliferative disease. Proc Natl Acad Sci USA. 2014;111:11467–72. doi:10.1073/pnas.1317751111.

39. Wang S, Wan X, Ruan Q. The MicroRNA-21 in Autoimmune Diseases. Int J Mol Sci. 2016;17. doi:10.3390/ijms17060864.

40. Swirski FK, Hilgendorf I, Robbins CS. From proliferation to proliferation: monocyte lineage comes full circle. Semin Immunopathol. 2014;36:137–48. doi:10.1007/s00281-013- 0409-1.

41. Lih-Brody L, Powell SR, Collier KP, Reddy GM, Cerchia R, Kahn E, et al. Increased oxidative stress and decreased antioxidant defenses in mucosa of inflammatory bowel disease. Dig Dis Sci. 1996;41:2078–86. doi:10.1007/BF02093613.

42. D’Incà R, Cardin R, Benazzato L, Angriman I, Martines D, Sturniolo GC. Oxidative DNA damage in the mucosa of ulcerative colitis increases with disease duration and dysplasia. Inflamm Bowel Dis. 2004;10:23–7. doi:10.1097/00054725-200401000-00003.

43. Dincer Y, Erzin Y, Himmetoglu S, Gunes KN, Bal K, Akcay T. Oxidative DNA damage and antioxidant activity in patients with inflammatory bowel disease. Dig Dis Sci. 2007;52:1636–41. doi:10.1007/s10620-006-9386-8.

44. Aslan M, Nazligul Y, Bolukbas C, Bolukbas FF, Horoz M, Dulger AC, et al. Peripheral lymphocyte DNA damage and oxidative stress in patients with ulcerative colitis. Pol Arch Med Wewn. 2011;121:223–9.

45. Beltrán B, Nos P, Dasí F, Iborra M, Bastida G, Martínez M, et al. Mitochondrial dysfunction, persistent oxidative damage, and catalase inhibition in immune cells of naïve and treated Crohn’s disease. Inflamm Bowel Dis. 2010;16:76–86. doi:10.1002/ibd.21027.

46. Liao J, Seril DN, Lu GG, Zhang M, Toyokuni S, Yang AL, et al. Increased susceptibility of chronic ulcerative colitis-induced carcinoma development in DNA repair enzyme Ogg1 deficient mice. Mol Carcinog. 2008;47:638–46. doi:10.1002/mc.20427.

47. Jones G-R, Bain CC, Fenton TM, Kelly A, Brown SL, Ivens AC, et al. Dynamics of colon monocyte and macrophage activation during colitis. Front Immunol. 2018;9:2764. doi:10.3389/fimmu.2018.02764.

48. Stagg AJ, Hart AL, Knight SC, Kamm MA. The dendritic cell: its role in intestinal inflammation and relationship with gut bacteria. Gut. 2003;52:1522–9.

49. Coats SR, Hashim A, Paramonov NA, To TT, Curtis MA, Darveau RP. Cardiolipins Act as a Selective Barrier to Toll-Like Receptor 4 Activation in the Intestine. Appl Environ Microbiol. 2016;82:4264–78. doi:10.1128/AEM.00463-16.

50. Levin A, Shibolet O. Toll-like receptors in inflammatory bowel disease-stepping into uncharted territory. World J Gastroenterol. 2008;14:5149–53. doi:10.3748/wjg.14.5149.

51. Horng T, Barton GM, Flavell RA, Medzhitov R. The adaptor molecule TIRAP provides signalling specificity for Toll-like receptors. Nature. 2002;420:329–33. doi:10.1038/nature01180.

52. Hug H, Mohajeri MH, La Fata G. Toll-Like Receptors: Regulators of the Immune Response in the Human Gut. Nutrients. 2018;10. doi:10.3390/nu10020203.

53. de Zoete MR, Bouwman LI, Keestra AM, van Putten JPM. Cleavage and activation of a Toll-like receptor by microbial proteases. Proc Natl Acad Sci USA. 2011;108:4968–73. doi:10.1073/pnas.1018135108.

54. Auffray C, Sieweke MH, Geissmann F. Blood monocytes: development, heterogeneity, and relationship with dendritic cells. Annu Rev Immunol. 2009;27:669–92. doi:10.1146/annurev.immunol.021908.132557.

