## Supplementary figures and images for "Extracellular vesicles produced by the human commensal gut bacterium *Bacteroides thetaiotaomicron* affect host immune pathways in a cell-type specific manner that are altered in inflammatory bowel disease"

### Supplementary Figure 1

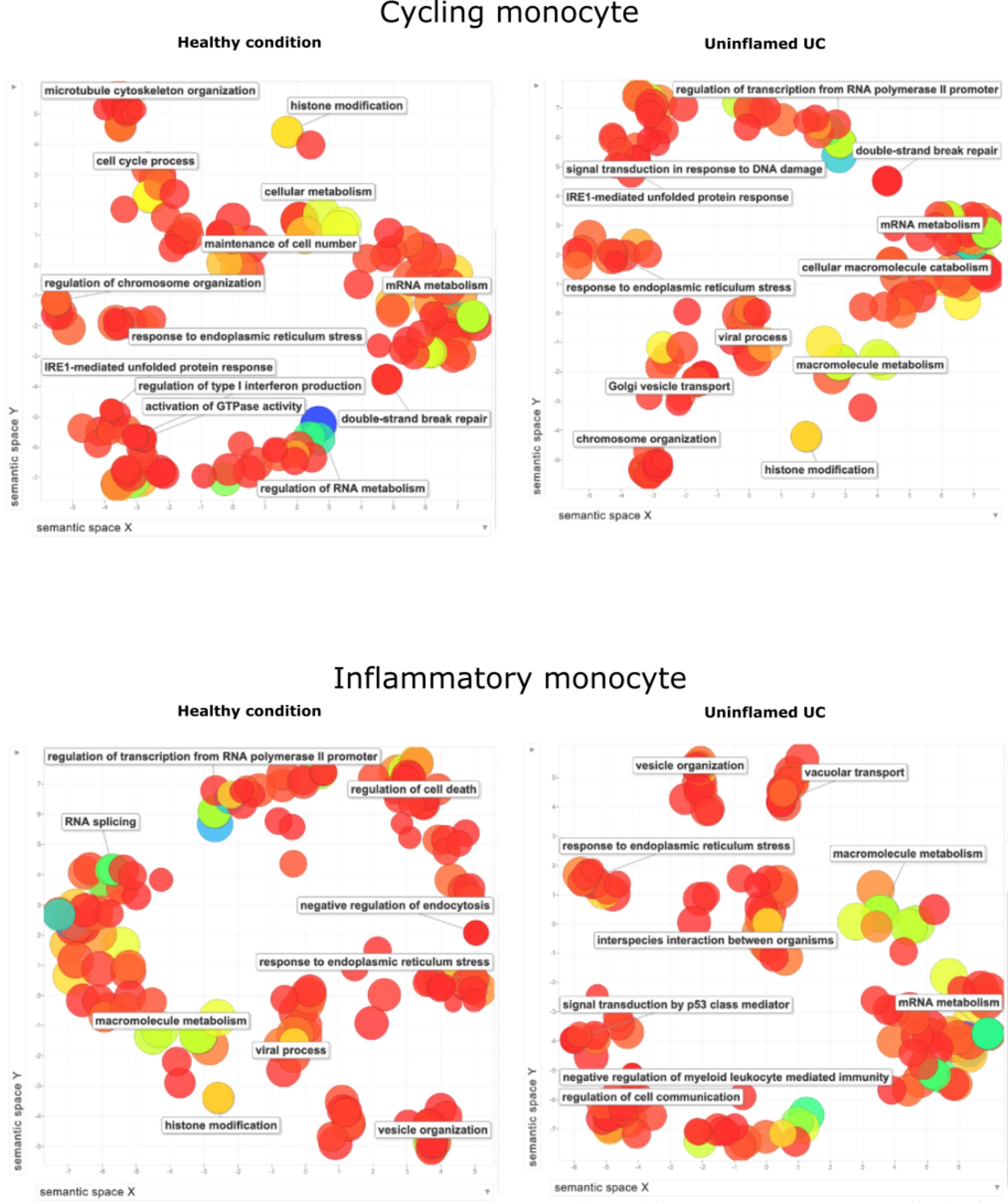

### Supplementary Figure 2

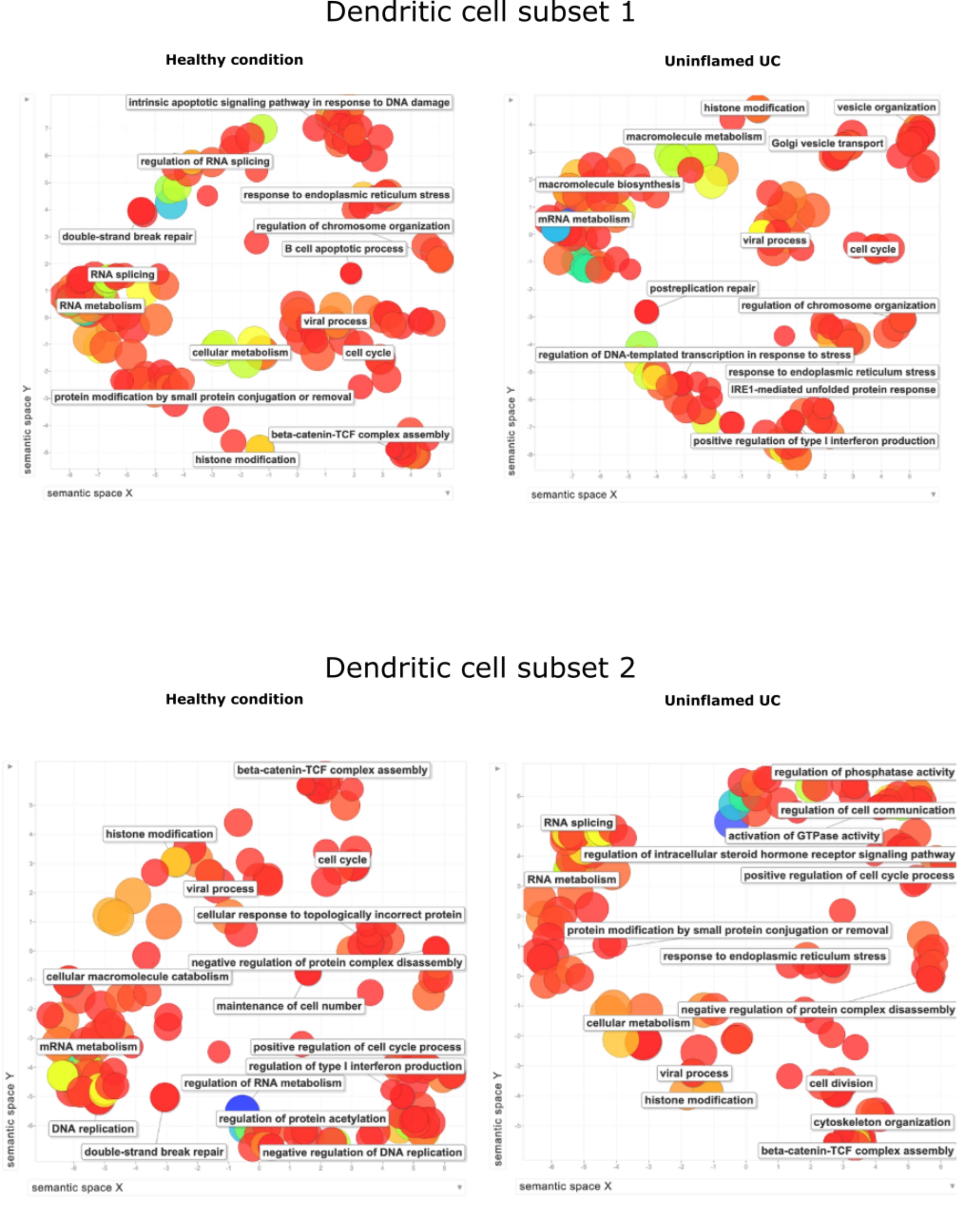

### Supplementary Figure 3

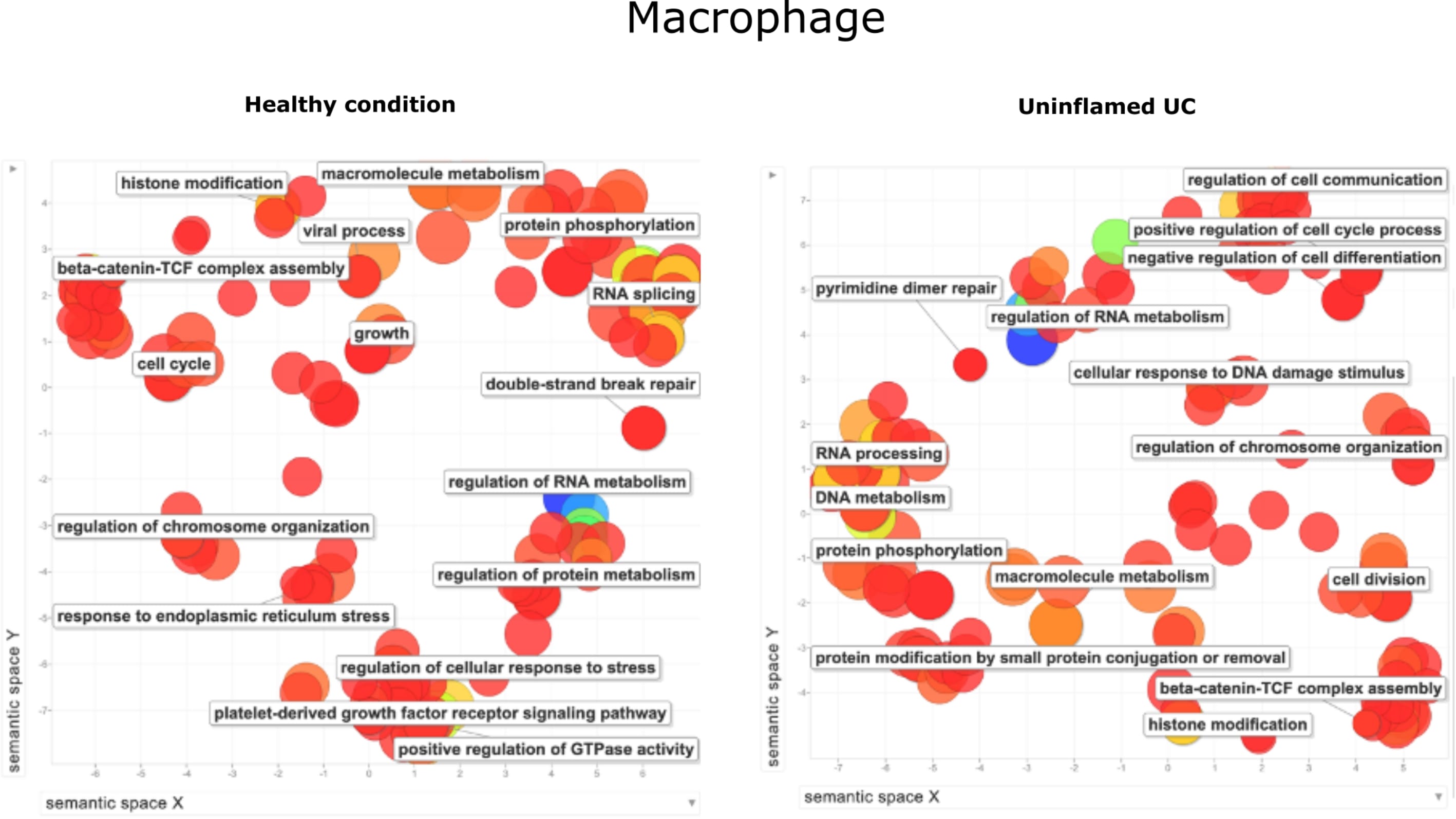
